# Spatial organization of *Clostridium difficile* S-layer biogenesis

**DOI:** 10.1101/405993

**Authors:** Peter Oatley, Joseph A. Kirk, Shuwen Ma, Simon Jones, Robert P. Fagan

## Abstract

Surface layers (S-layers) are protective protein coats which form around all archaea and most bacterial cells. *Clostridium difficile* is a Gram-positive bacterium with an S-layer covering its peptidoglycan cell wall. The S-layer in *C. difficile* is constructed mainly of S-layer protein A (SlpA), which is a key virulence factor and an absolute requirement for disease. S-layer biogenesis is a complex multi-step process, disruption of which has severe consequences for the bacterium. We examined the subcellular localization of SlpA secretion and S-layer growth; observing formation of S-layer at specific sites that coincide with cell wall synthesis, while the secretion of SlpA from the cell is relatively delocalized. We conclude that this delocalized secretion of SlpA leads to a pool of precursor in the cell wall which is available to repair openings in the S-layer formed during cell growth or following damage.

## Introduction

*Clostridium difficile* infection (CDI) is the major cause of antibiotic associated diarrhoea (Hull & Beck, 2004) and can lead to severe inflammatory complications (Napolitano & Edmiston, 2017). This Gram-positive bacterium has a cell wall encapsulating, proteinaceous surface-layer (S-layer), a paracrystalline array that acts as a protective semipermeable shell and is essential for virulence (Kirk et al., 2017). In *C. difficile* the S-layer consists mainly of SlpA, the most abundant surface protein (Calabi et al., 2001). SlpA is produced as a pre-protein (Figure 1A) that is secreted and processed by the cell wall cysteine protease Cwp84 into low molecular weight (LMW) and high molecular weight (HMW) SLP subunits (Kirby et al., 2009)(Figure 1B). These two subunits form a heterodimeric complex that is then incorporated into the crystalline lattice of the S-layer, which is anchored to cell wall polysaccharide PS-II via three cell wall binding (CWB2) motifs within the HMW region (Fagan et al., 2009; Willing et al., 2015) (Figure 1A).

**Figure 1:**
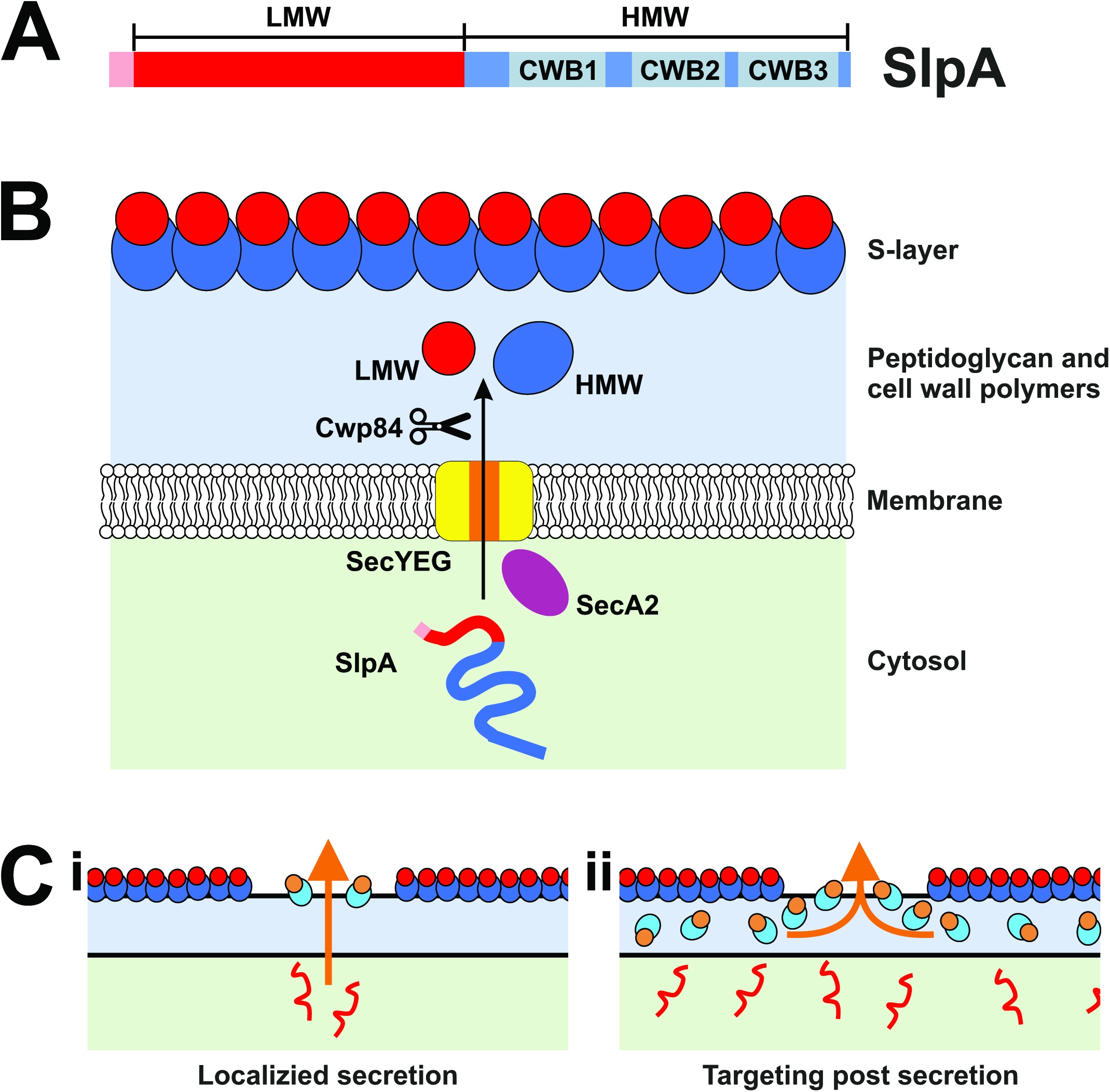
The *C. difficile* S-layer and the SlpA secretory pathway. **A:** Domain structure of SlpA precursor protein with signal sequence (Pink), low molecular weight region (LMW, Red) and high molecular weight region (HMW, Blue) that contains three cell wall binding domains (CWB, 1-3 in Grey). **B:** Schematic diagram of SlpA secretion and processing in *C. difficile*. SlpA (Pink/Red/blue line) is translated in the cytosol (light green) and targeted for secretion across the membrane using SecA2 (Purple oval) most likely via the SecYEG Channel (Yellow/Orange). Cwp84 (Scissors) cleaves SlpA into low molecular weight (LMW, Red spheres) and high molecular weight (HMW, Blue spheres) S-layer protein (SLP) subunits. The HMW and LMW SLPs assemble to form hetero-dimers that incorporate into the S-layer. The surface of the S-layer consists largely of exposed LMW-SLP anchored to the cell wall (light blue) via cell wall biding domains of the HMW-SLP component (see A). **C:** Models of SlpA integration into the S-layer. (i) unfolded SlpA (red line) is secreted from specific points on the cell membrane - directed by gaps in the S-layer or cell wall (colored as in B), newly processed SlpA (LMW-SLP orange circles, HMW-SLP light blue ovals) is transported directly through the cell wall for integration into the S-layer. Alternatively; (ii) SlpA is translocated across the cell membrane at multiple sites. A pool of SlpA lays within the cell wall ready to fill gaps in the S-layer.

The production and secretion of S-layer components are energetically expensive for the cell, suggesting that the process will display evolved efficiency. However, it is not yet clear how S-layer formation is spatially regulated and whether SlpA is targeted to areas of cellular growth before or after secretion (Figure 1C). *C. difficile* express two homologs of the *E. coli* cytosolic protein export ATPase, SecA: SecA1 and SecA2 (Fagan & Fairweather, 2011). These two SecAs are thought to promote post-translational secretion through the general secretory (Sec) pathway. SecA2 is required for efficient SlpA secretion (Fagan & Fairweather, 2011) and is encoded adjacent to *slpA* on the chromosome (Monot et al., 2011). It has been shown that some SecA2 systems secrete specific substrates (reviewed by (Bensing, Seepersaud, Yen, & Sullam, 2014)) which may ease the burden on the general Sec system and allow spatial or temporal regulation of secretion.

As an obligate anaerobe, *C. difficile* has been notoriously difficult to visualize using standard microscopy techniques with commonly used oxygen-dependent fluorescent proteins and this is further complicated by intrinsic autofluorescence in the green spectrum (Ransom, Ellermeier, & Weiss, 2015). To circumvent these problems, we have used a variety of labeling techniques to avoid the requirement for oxygen maturation and any overlap with autofluorescence. Using fluorescence microscopy, we identified areas of S-layer biogenesis and SlpA secretion to determine if this S-layer component is specifically targeted to growing parts of the cell. Firstly, we probed the localization of newly synthesized S-layer which was detected at discrete regions which coincided with areas of new cell wall biosynthesis. We continued by studying the internal localization of SecA2 and SlpA, discovering that SlpA is secreted all over the cytoplasmic membrane. Having observed delocalized secretion of SlpA, yet localized new surface S-layer, we conclude that there is a pool of SlpA that resides within the cell wall which is available to construct regions of the developing S-layer.

## Results

### Newly synthesized S-layer co-localizes with areas of new cell wall

During exponential growth, *C. difficile* cells are constantly growing and dividing, requiring the production of new peptidoglycan at the cell wall. The S-layer protects the cell envelope from innate immune effectors such as lysozyme and LL-37 (Kirk et al., 2017). This function requires that an S-layer barrier is maintained while new peptidoglycan is synthesized during growth. Peptidoglycan can be labelled by growing *C. difficile* cells in the presence of the fluorescent D-amino acid, HCC-amino-D-alanine (HADA), (Kuru et al., 2012). Subsequent chasing with unlabeled media and imaging of live cells (Figure 2A and Video 1) or fixed cells over a time course (Figure 2-figure supplement 1) reveals sites of newly synthesized peptidoglycan that appear less intense for HADA. This pattern of HADA staining is seen at the dividing septum and along the long axis of the cell (Figure 2). Combining this with the inducible expression of the immunologically distinct SlpA_R20291_ in *C. difficile* strain 630 (Figure 2-figure supplement 2A), allowed areas of newly assembled S-layer to be visualized by immunofluorescence (Figure 2B and Figure 2-figure supplement 2B). Tracing the intensity of cell wall staining with the signal from newly synthesized surface SlpA reveals a crude anti-correlation (Figure 2C and Figure 2-figure supplement 3) and suggests that new S-layer is formed at areas of newly formed underlying cell wall. The areas of newly synthesized cell wall that are void of SlpA_R20291_ signal are likely to be filled with endogenous SlpA_630_ that is expressed at much higher levels, as observed in extracellular cell wall protein extracts (Figure 2-figure supplement 2A).

**Figure 2:**
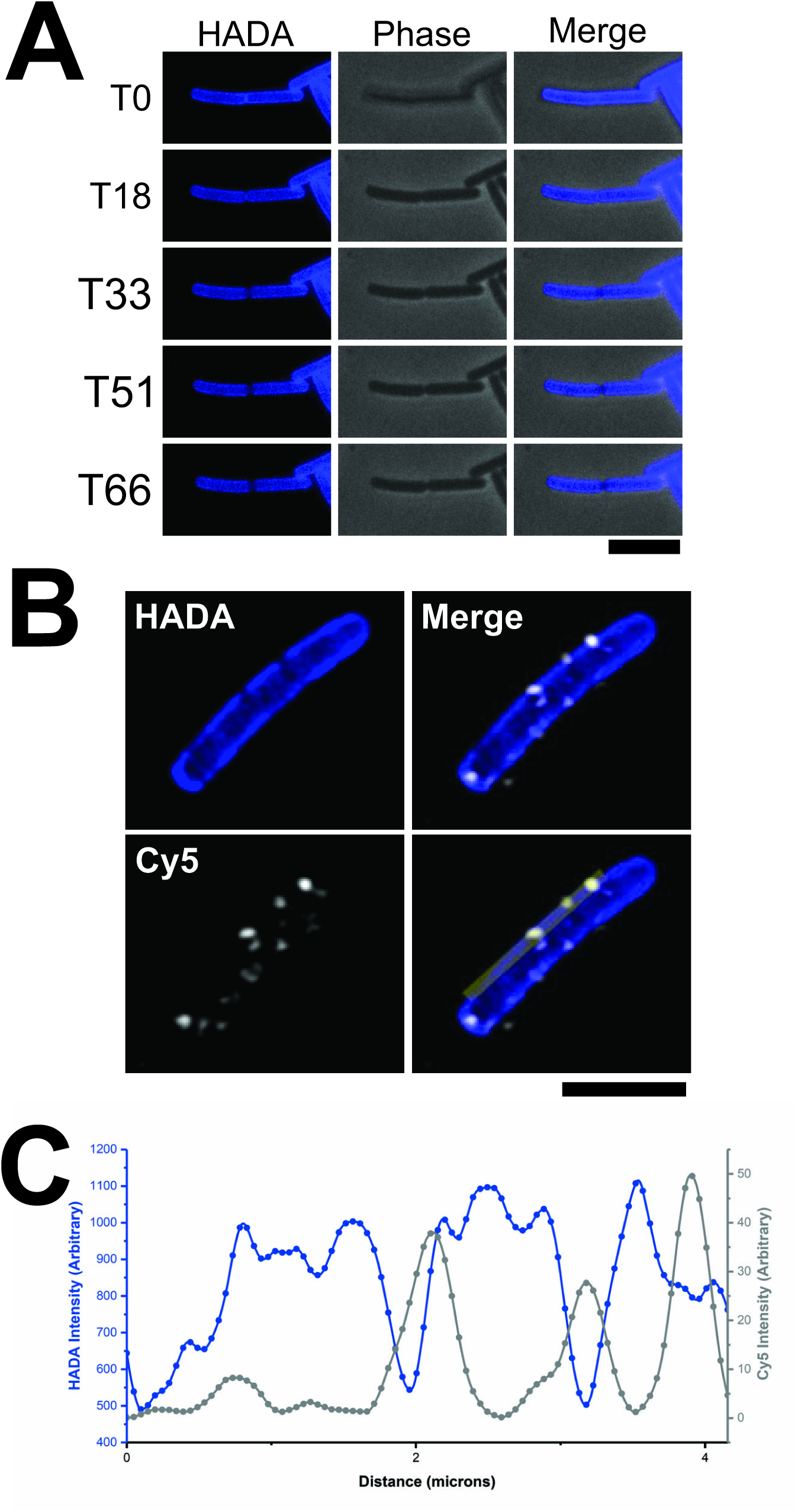
New surface S-layer colocalizes with areas of new peptidoglycan synthesis. **A:** Examples of timepoints from real-time widefield fluorescent HADA signal (left panels) and phase contrast (center panels) of *C. difficile* 630 cells chased for HADA stain. Frame time represented in minutes, scale bar indicates 6 µm. **B:** Airyscan confocal image of a *C. difficile* 630 cell grown with HADA to label peptidoglycan cell wall (Blue) and chased to reveal darker areas of newly synthesized cell wall. This chase was followed by a short expression of SlpA_R20291_ which was specifically immunolabeled with Cy5 (White). Yellow bar indicates the region used for the intensity plot in C. Scale bar indicates 6 µm. **C:** Intensity plot depicting signal from HADA (Blue) and Cy5 (Grey) along the yellow bar illustrated in B.

During cell division, a large amount of new surface SlpA can be detected at the septum (Figure 3). This staining pattern suggests that S-layer is actively formed on the mother cell over the newly synthesized cell wall, preparing each daughter cell with a new S-layer cap before cell division is complete. Numerous cells display areas of new cell wall at one of their poles that co-insides with new S-layer staining (Figure 3). We interpret these as new daughter cells that have completed cell division during the HADA stain chase as detected in live cell imaging (Figure 2A and Video 1). New polar S-layer can be sorted into three categories: staining distributed over the whole cell cap, on the tip of the cap or at the sides of the new cap close to the older cell wall (Figure 3). Daughter cells with their poles completely covered in new S-layer have most likely expressed SlpA_R20291_ throughout cell division and have SlpA_R20291_ distributed all over the new S-layer cap. The apex of the cell cap marks the final place of new daughter cell formation and those caps stained just at the tip have probably expressed SlpA_R20291_ towards the final stages of division as the two daughter cells separate and the cap is completed. Areas stained at the connecting edge between the pole and the main body of the cell must represent areas of growth once the S-layer cap was fully formed when cell division was completed. Together, these staining patterns support the hypothesis that S-layer is assembled on the mother cell at the septum to form polar caps for the daughter cells to maintain a continuous protective barrier following cell separation.

**Figure 3:**
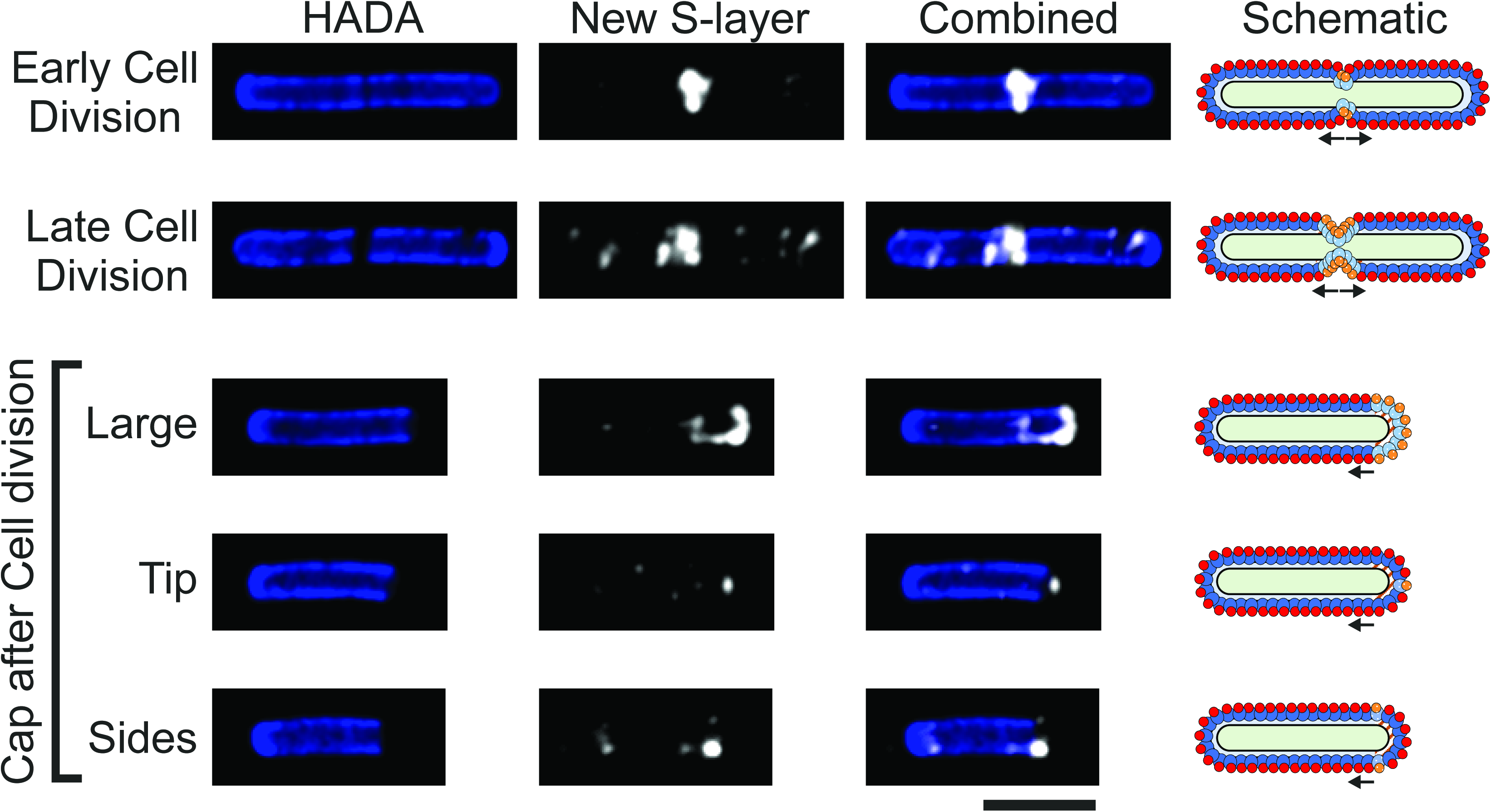
S-layer formation during cell division. Airyscan confocal images of *C. difficile* 630 cells during and immediately after cell division with HADA labelled peptidoglycan cell wall (blue) and new surface SlpA_R20291_ immunolabeled with Cy5 (white). Large, dark areas lacking HADA staining mark cell wall synthesis at the septum between cells or a newly produced cell pole. Scale bar indicates 3 µm. On the right-hand side of each row is a schematic diagram illustrating the position of new surface SlpA_R20291_ (HMW-SLP, spotted light blue and LMW-SLP, spotted orange) as detected in the corresponding microscopy images against the position of endogenous surface SlpA_630_ (HMW-SLP, dark blue and LMW-SLP, red). The position of newly synthesized cell wall is displayed in brown/white stripes.

### SecA2 localization

As newly synthesized S-layer is formed at specific points on the cell surface (Figure 3) we wanted to determine if these areas correlate with concentrated points of SlpA secretion from the cytosol. Having designated sites of secretion would allow the efficient targeting of S-layer precursors to where they are needed. As it has been shown that SecA2 is essential for cell survival (Dembek et al., 2015) and performs a critical role in SlpA secretion (Fagan & Fairweather, 2011), we assumed that intracellular positioning of SecA2 will reveal where SlpA is secreted. To confirm this, we set out to create a functional, fluorescently tagged SecA2 for monitoring SecA2 localization by microscopy. A *C. difficile* strain 630 mutant was generated that encodes a C-terminal SNAP-tagged SecA2 (SecA2-SNAP) on the genome in the original locus and under the control of the native promoter. SecA2-SNAP was the only SecA2 protein detected in membrane fractions by western immunoblot analysis (Figure 4-figure supplement 1A) and, when stained with TMR-Star, this protein species was the only one visualized by in-gel fluorescence (Figure 4-figure supplement 1B). Cells expressing SecA2-SNAP displayed similar growth dynamics to the wild-type parental strain (Figure 4-figure supplement 1C), suggesting that the fusion protein is fully functional as SecA2 is essential for growth (Dembek et al., 2015). Imaging cells by widefield microscopy revealed that SecA2 is distributed throughout the cell and not localized to specific areas (Figure 4A). Higher resolution, Airyscan confocal images revealed the same widespread distribution but with pockets of higher intensity signal (Figure 4B). By combining SecA2 localization with HADA chase staining and new S-layer labelling, no correlation between SecA2 within the cell and areas of newly synthesized S-layer on the cell periphery could be identified (Figure 4B). Together these data suggest that SecA2 is not the determining factor in localization of new S-layer growth.

**Figure 4:**
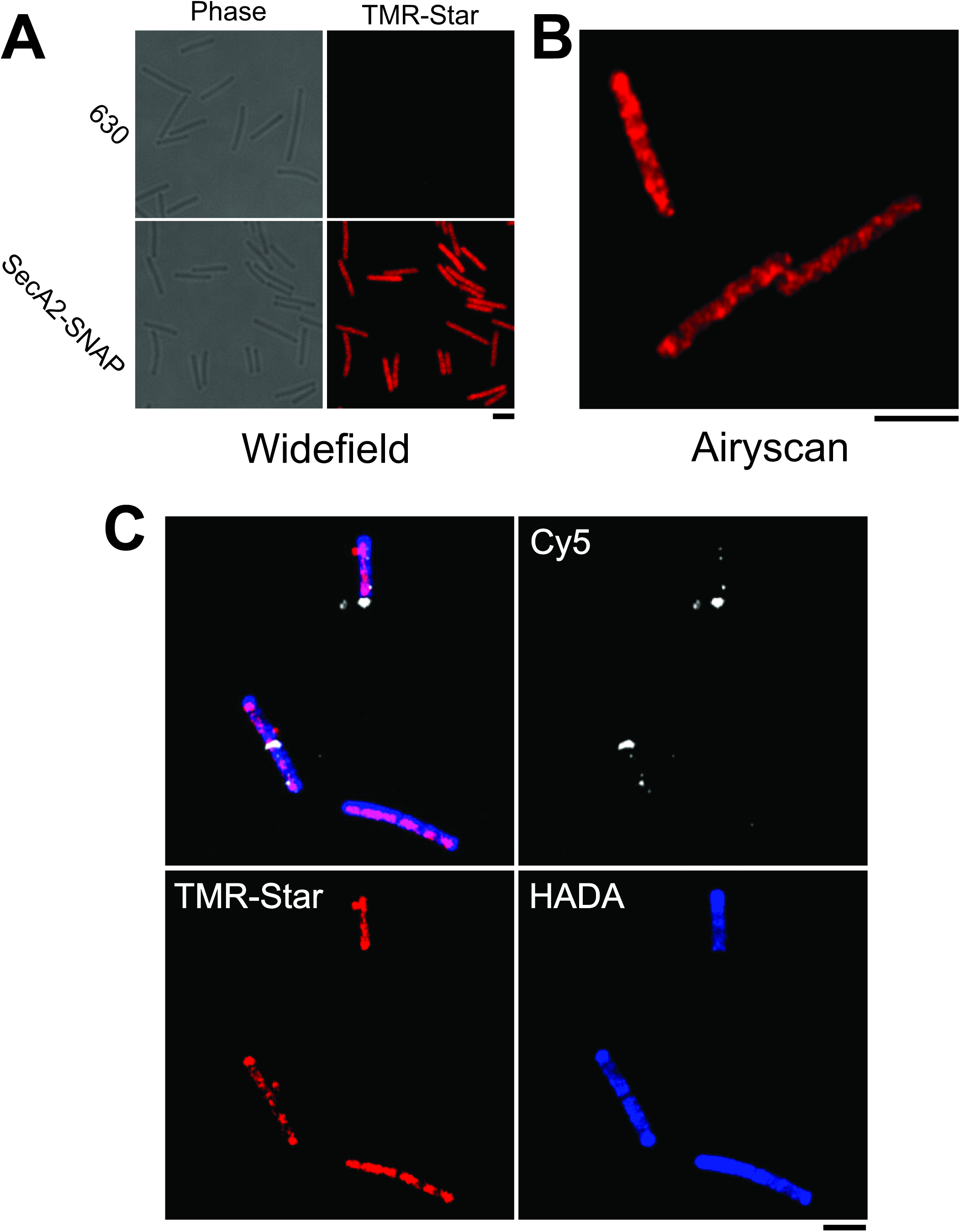
SecA2-SNAP localization and new S-layer. **A:** Widefield phase contrast (left panels) and fluorescent (right panels) images of wild type *C. difficile* 630 or 630 *secA2-snap* cells stained with TMR-Star (red). Scale bar indicates 3 µm. **B:** Airyscan confocal image displaying SecA2-SNAP-TMR-Star signal distribution in *C. difficile* 630 cells. Scale bar indicates 3 µm. **C:** Airyscan confocal image showing the localization of SecA2-SNAP-TMR-Star (red) in relation to the synthesis of cell wall (dark patches lacking blue HADA stain) and newly synthesized S-layer (Cy5, white). Scale bar indicates 3 μm.

### SlpA secretion

Although SecA2 was visualized throughout the cell, SecA2 has additional secretory substrates (Fagan et al., 2011) so it is possible that secretion of SlpA itself may be localized. Although immunofluorescence has been used to detect surface SlpA, S-layer pore size is thought to be too small to allow the access of antibodies to proteins located in the cell wall or indeed within the cell (Fagan & Fairweather, 2014). To determine where SlpA is being secreted, two different SlpA fusion proteins were constructed, an SlpA-SNAP fusion that can be secreted and is found in the extracellular fraction and a SNAP tagged SlpA-dihydrofolate reductase (SlpA-DHFR-SNAP) fusion that associates with the cellular fraction (Figure 5A & Figure 5-figure supplement 1A). DHFR is a fast folding protein that has been used to block and probe protein translocation mechanisms for many years (Arkowitz, Joly, & Wickner, 1993; Bonardi et al., 2011; Eilers & Schatz, 1986; Rassow et al., 1989). Expressing SlpA-DHFR-SNAP decreases the secretion of native *C. difficile* extracellular proteins (Figure 5-figure supplement 1B) and leads to the build-up of precursor SlpA within the cell (Figure 5-figure supplement 2), consistent with DHFR blocking the SecA2 translocon. This effect requires the SlpA signal sequence, as an SlpA-DHFR-SNAP lacking the N-terminal signal sequence no longer blocks protein translocation (Figure 5-figure supplement 1B). Together these findings show that the SlpA-DHFR fusion used here is specifically targeted to and occupies the same secretory channel required for wild-type SlpA secretion. To prevent protein secretion, the DHFR domain must first fold correctly (Arkowitz et al., 1993) and will therefore only prevent secretion via the post-translational pathway. The lack of detectable SlpA-DHFR-SNAP in the extracellular fraction (Figure 5-figure supplement 1A) shows for the first time that SlpA is exclusively post-translationally translocated. Using these two SNAP fusion proteins we can now probe the intercellular localization of SlpA secretion (SlpA-DHFR-SNAP) and localization once secreted (SlpA-SNAP).

**Figure 5:**
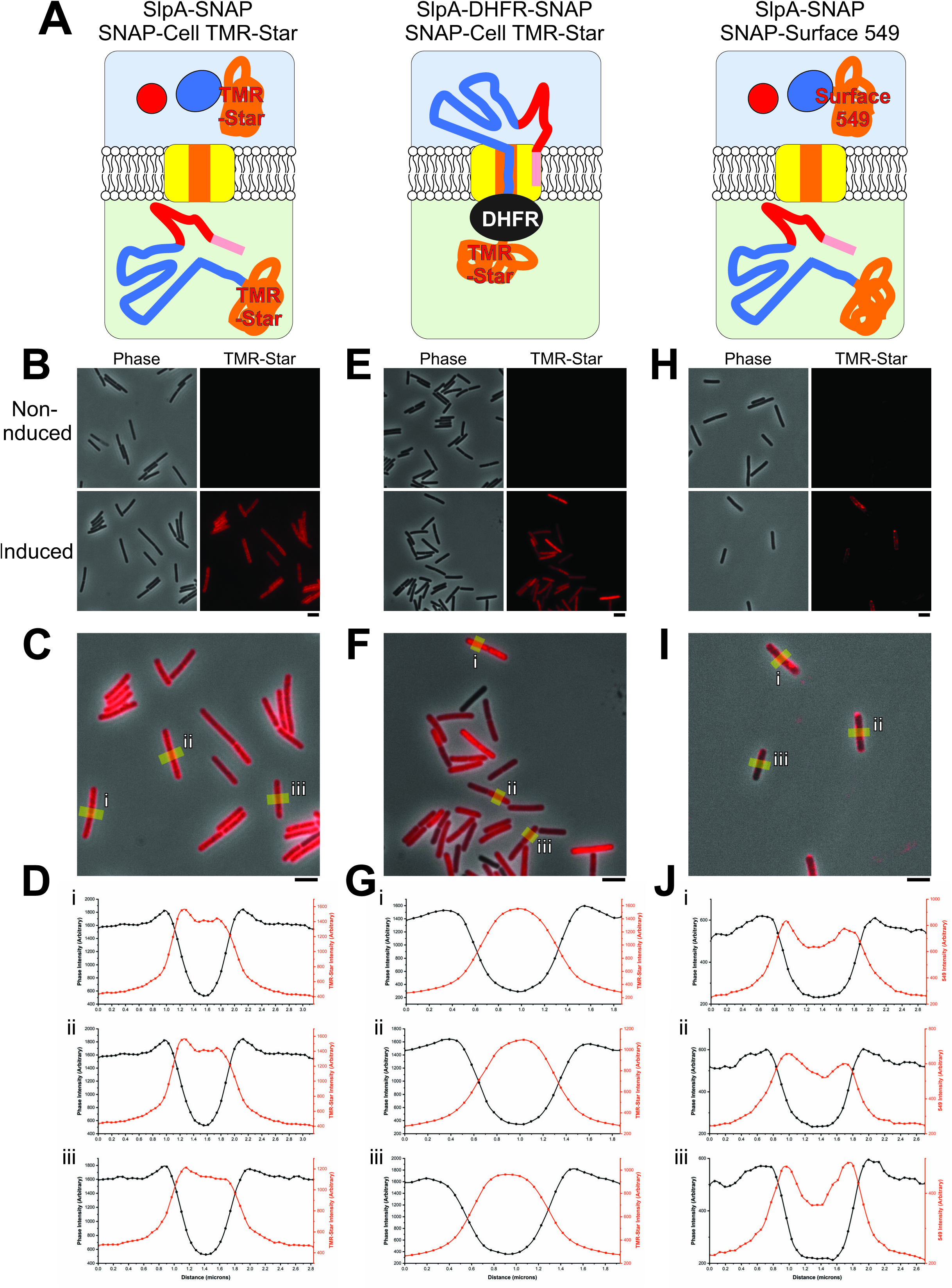
Sites of S-layer secretion. **A:** Schematic diagram illustrating the position of stained SNAP tagged SlpA constructs expressed in *C. difficile* 630 cells: SNAP-Cell TMR-Star stained SlpA-SNAP (left panel) or SlpA-DHFR-SNAP (center panel) and SNAP-Surface 549 stained SlpA-SNAP (right panel). Colored as in Figure 1B with SNAP tags represented as an orange coil. SlpA-SNAP is exported and cleaved into LMW-SLP and HMW-SLP-SNAP. The DHFR domain (dark gray oval) of SlpA-DHFR-SNAP blocks the translocon channel during export, leaving the TMR-Star bound SNAP tag in the cytosol. SNAP-Surface 549 stains extracellular HMW-SLP-SNAP only. **B:** Widefield phase contrast (left panels) and fluorescent SNAP-Cell TMR-Star signal (right panels) of *C. difficile* 630 cells stained with SNAP-Cell TMR-Star imaged with and without induction of SlpA-SNAP expression. Scale bar indicates 3 µm. **C:** Overlay of fluorescent signal in the induced sample (from B) with areas taken for the plot profiles labelled (yellow lines, i-iii). Scale bar indicates 3 µm. **D**: SlpA-SNAP-Cell TMR-Star profile plots of i-iii (from C) of phase contrast signal (black) and SNAP-Cell TMR-Star signal (red). **E:** SlpA-DHFR-SNAP in *C. difficile* 630 cells (labelled as in B). **F**: Overlay of signal in the induced sample (from E, labelled as in C). **G**: SlpA-DHFR-SNAP-Cell TMR-Star profile plots of i-iii (from F) (labelled as in D). **H:** Widefield phase contrast (left panels) and fluorescent SNAP-Surface 549 signal (right panels) of *C. difficile* 630 cells stained with SNAP-Surface 549 imaged with and without induction of SlpA-SNAP expression. **I:** Overlay of signal in the induced sample (from H, labelled as in C). **J:** HMW-SLP-SNAP-Surface 549 profile plots of i-iii (from I) of phase contrast signal (black) and SNAP-Surface 549 (red).

After secretion, the SlpA-SNAP protein is processed as normal by the cell wall localized cysteine protease Cwp84, yielding the LMW-SLP subunit and a SNAP tagged HMW-SLP (HMW-SLP-SNAP). Widefield images show a diffuse distribution of HMW-SLP-SNAP on the cell surface with a halo of TMR-Star signal surrounding most of the cells (Figure 5B and C). Surface intensity plots reveal a broad cross-section of TMR-Star intensity across the cell width (Phase vs TMR-Star signal, Figure 5D). Within these cross-sections there are smaller peaks in TMR-Star intensity which correlate with the cell periphery, displayed as rapid changes in phase signal (Figure 5D), this is consistent with the detection of HMW-SLP-SNAP TMR-Star in the extracellular fraction (Figure 5-figure supplement 1A). Cells expressing SlpA-DHFR-SNAP show some heterogeneity of expression (Figure 5E and F), perhaps caused by leaky expression leading to a negative selective pressure for the plasmid and the drastic effects this DHFR fusion protein has on secretion (Figure 5-figure supplement 1C). Again, the signal from SlpA-DHFR-SNAP appears diffuse throughout the cell and not located at specific sites (Figure 5E and F). Signal intensity traces reveal a narrow TMR-Star signal peak towards the interior of the cell where the phase signal is low (Figure 5G), suggesting a more intracellular location for SlpA-DHFR-SNAP than HMW-SLP-SNAP which is consistent with SlpA-DHFR-SNAP being trapped in the cell at the membrane (Figure 5-figure supplement 1A).

To obtain a clearer image of newly secreted extracellular HMW-SLP-SNAP, cells were treated with SNAP-Surface 549 (Figure 5H and I) that specifically stains extracellular proteins (Figure 5-figure supplement 1A) and should reduce the intracellular SlpA-SNAP background staining observed with SNAP-Cell TMR-Star (Figure 5A and Figure 5-figure supplement 1A). HMW-SLP-SNAP-Surface 549 outlines of cells were visible in widefield (Figure 5H) and Airyscan confocal imaging (Figure 5-figure supplement 3). Surface density plots of cross-sections of these cells have SNAP-Surface 549 signal peaks towards the cell periphery (Figure 5J) which supports the similar extracellular staining pattern seen with SNAP-cell TMR-star (Figure 5D). However, the intensity of the HMW-SLP-SNAP-Surface 549 appears uneven and pockets of higher intensity signal can be observed (Figure 5-figure supplement 3) that may relate to where HMW-SLP accumulates post-secretion and where this protein inserts into the S-layer. The distribution of signal in these images suggest that SlpA secretion occurs over the majority of the cell’s surface and not just at sites where new S-layer is being formed.

## Discussion

For an S-layer to function correctly it must completely encapsulate the cell (de la Riva, Willing, Tate, & Fairweather, 2011; Kirk et al., 2017). We propose here that S-layer is assembled at areas of newly synthesized peptidoglycan to maintain a stable S-layer that continually protects the *C. difficile* cell. Although newly synthesized SlpA is secreted from all regions of the cell, only a relatively small proportion of this was detected at the surface. This irregularity suggests that *C. difficile* possess reserves of SlpA beneath the S-layer in the cell wall (Figure 6). Although excess SlpA production and storage will be quite energetically expensive for the cell, this reservoir of SlpA could provide a positive fitness advantage by allowing cells to respond quickly to repair gaps in this critical barrier (Figure 6). Examples of self-repair mechanisms are present thorough all forms of life from intracellular vesicular mediated membrane healing (McNeil & Baker, 2001; Tang & Marshall, 2017) up to a tissue level such as wound healing (Greaves, Ashcroft, Baguneid, & Bayat, 2013). In addition to allowing rapid repair, by having a stockpile of SlpA in the cell wall, *C. difficile* may also create a buffer to reduce the amount of *de novo* SlpA translation and translocation required to safely complete cell division. It is not clear how large this buffer is and it would be interesting to identify the proportion of SlpA that lays below the S-layer, in reserve.

**Figure 6:**
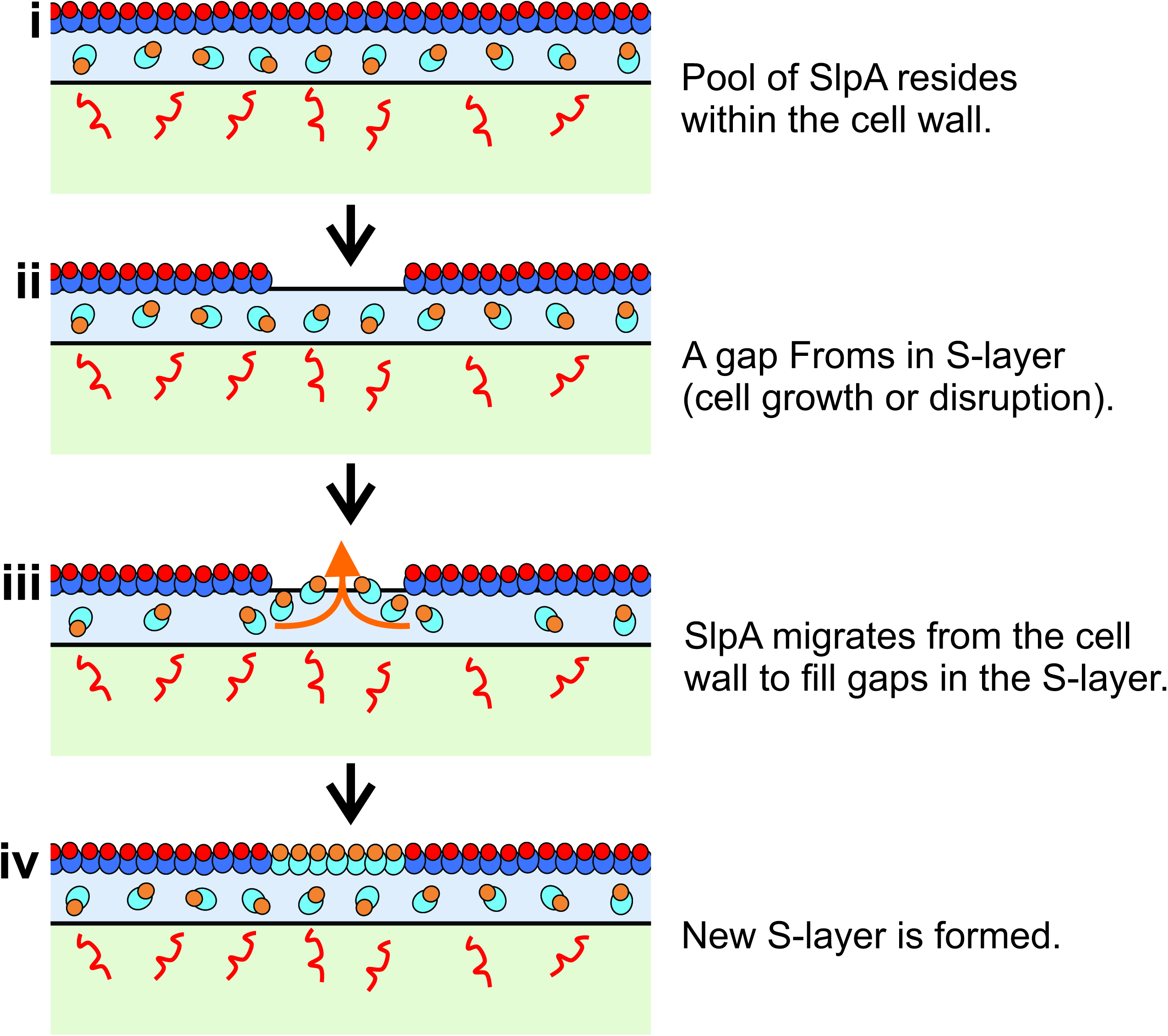
Model of S-layer in the cell wall. Schematic flow diagram of SlpA secretion and S-layer formation (colored as in Figure 1C). During normal cell growth SlpA is targeted by SecA2 for secretion all over the cytosolic membrane. A store of SlpA resides within the cell wall where it is processed ready for integration into the S-layer (i). Gaps may form in the S-layer due to cell growth or injury (ii). SlpA in the cell wall diffuses out (iii) and fills openings in the S-layer (iv).

Although the S-layer is a rigid structure (Mescher & Strominger, 1976), fractures in the S-layer must form to allow the cells to grow. Our data suggests that these fractures coincide with new peptidoglycan synthesis and that new SlpA emerges through these gaps to be incorporated into the crystalline lattice. Higher resolution imaging techniques may allow the direct observation these gaps in the S-layer and how the separate S-layer sections assemble. When new S-layer and secreted HMW-SLP-SNAP was labelled (Figure 2B, Figure 2-figure supplement 3 and Figure 5-figure supplement 3) and intracellular SecA2-SNAP was detected (Figure 4B), regular patterns and sometimes diagonal staining could be seen along the longitudinal axis of the cell. These patters may relate the localization of SecA2 and newly forming S-layer in line with intracellular cytoskeletal and motor proteins that power cell growth (Colavin, Shi, & Huang, 2018).

We have also demonstrated for the first time that SlpA is secreted post-translationally. Proteins transported in this way usually interact with cytosolic chaperones that prevent folding prior to translocation (Kim & Kendall, 2000). The identity of these chaperones and the exact role SecA2 plays in SlpA secretion has yet to be determined. Since SlpA must undergo a post-secretion protease modification (Figure 1) (Kirby et al., 2009), having a dwell time in the cell wall will allow time for correct processing. However this also poses the question of how S-layer components located there are prevented from oligomerizing (Takumi, Koga, Oka, & Endo, 1991) prior to assembly at the surface. It is tempting to speculate that the S-layer assembly pathway may also involve extracellular chaperones to facilitate processing and targeting while preventing premature self-assembly. The revelation that there is a pool of SlpA in the cell wall and the accessibility of the cell wall to drugs may provide opportunities for the identification of novel narrow spectrum targets that affect the assembly of this essential virulence factor.

In summary, we have found that S-layer is formed at sites of cell wall synthesis and there is an underlying supply of the S-layer precursor, SlpA, located throughout the cell wall.

## Methods

### Media and Growth Conditions

All strains, plasmids and oligonucleotides used in this investigation are displayed in Table 1. CA434 and NEB5α *E. coli* were grown in LB broth or on LB agar supplemented when required with 15 µg/ml chloramphenicol for plasmid selection. *C. difficile* were grown in reduced TY (3% tryptose, 2% yeast extract) broth or on Brain Heart Infusion agar under strict anaerobic conditions. Cultures were supplemented with 15 µg/ml thiamphenicol when selecting for plasmids.

**Table 1:**
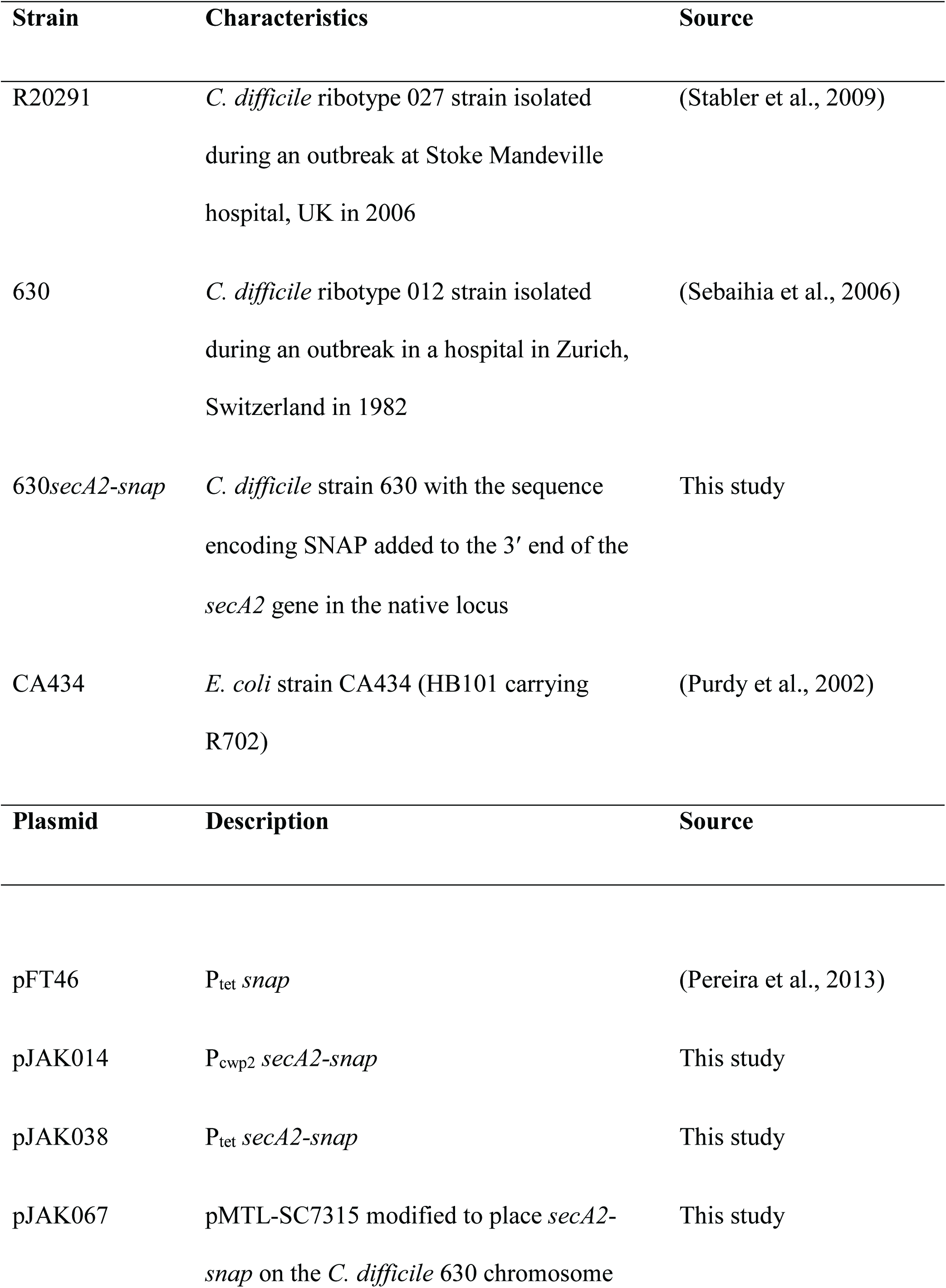
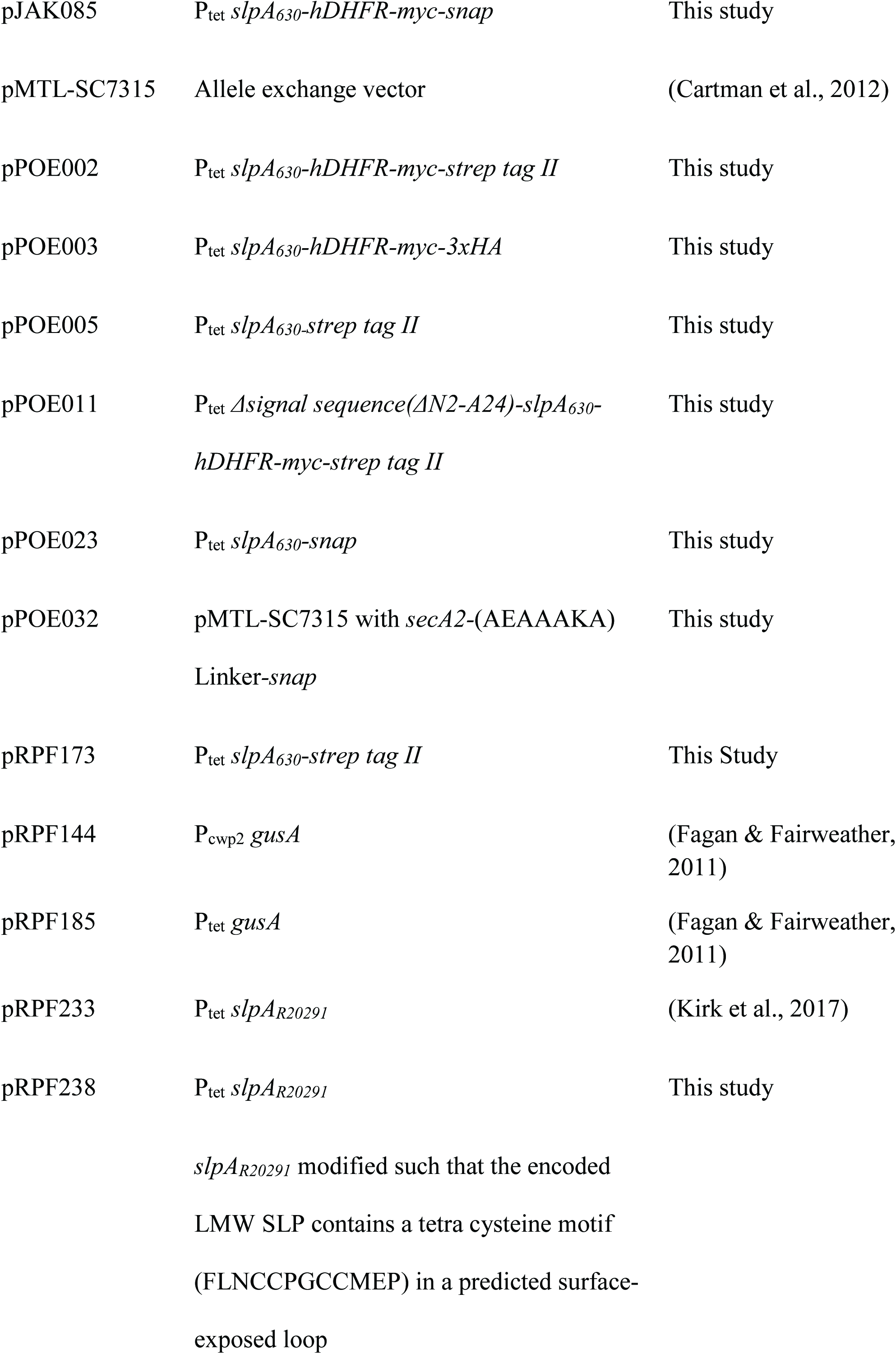
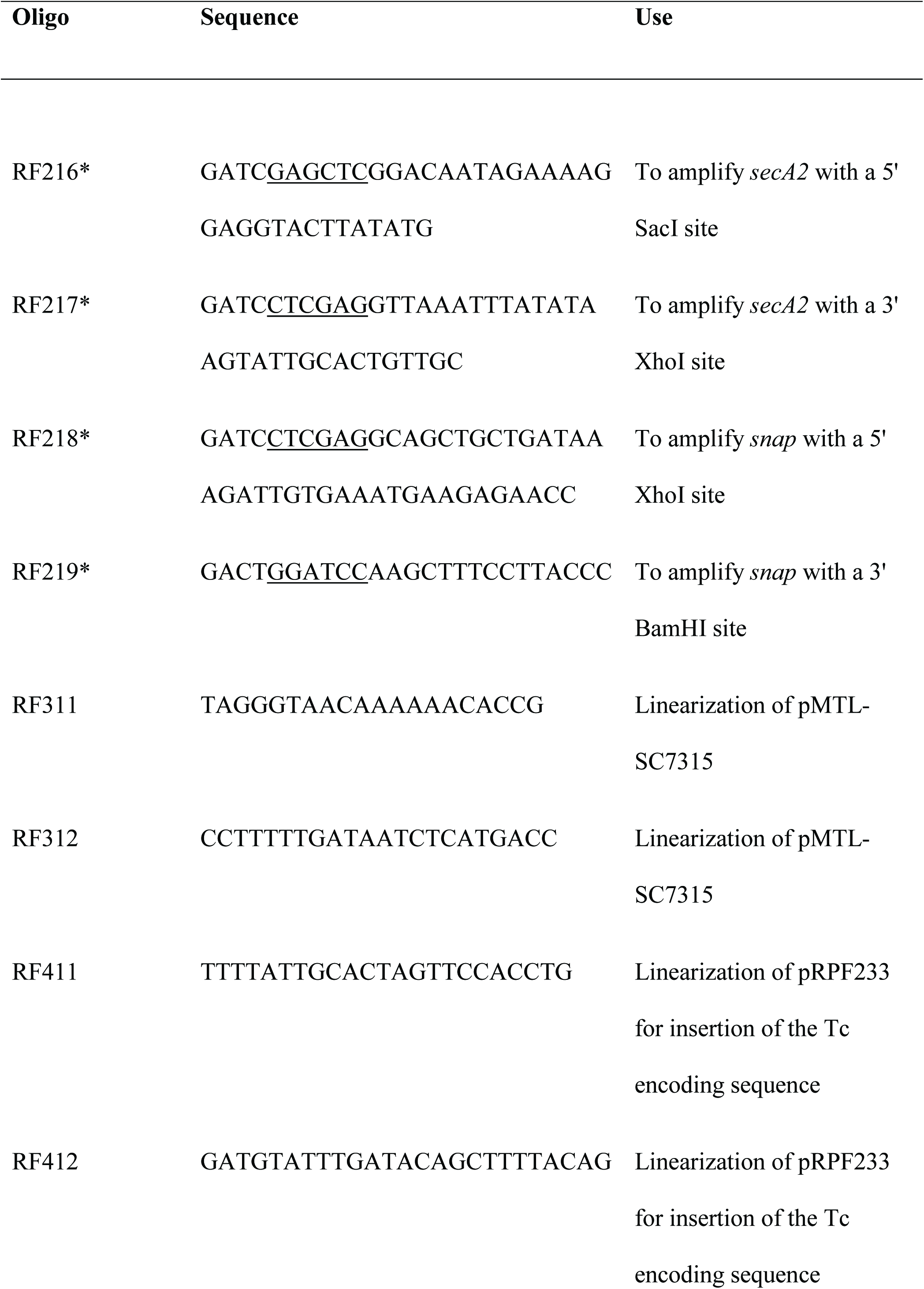
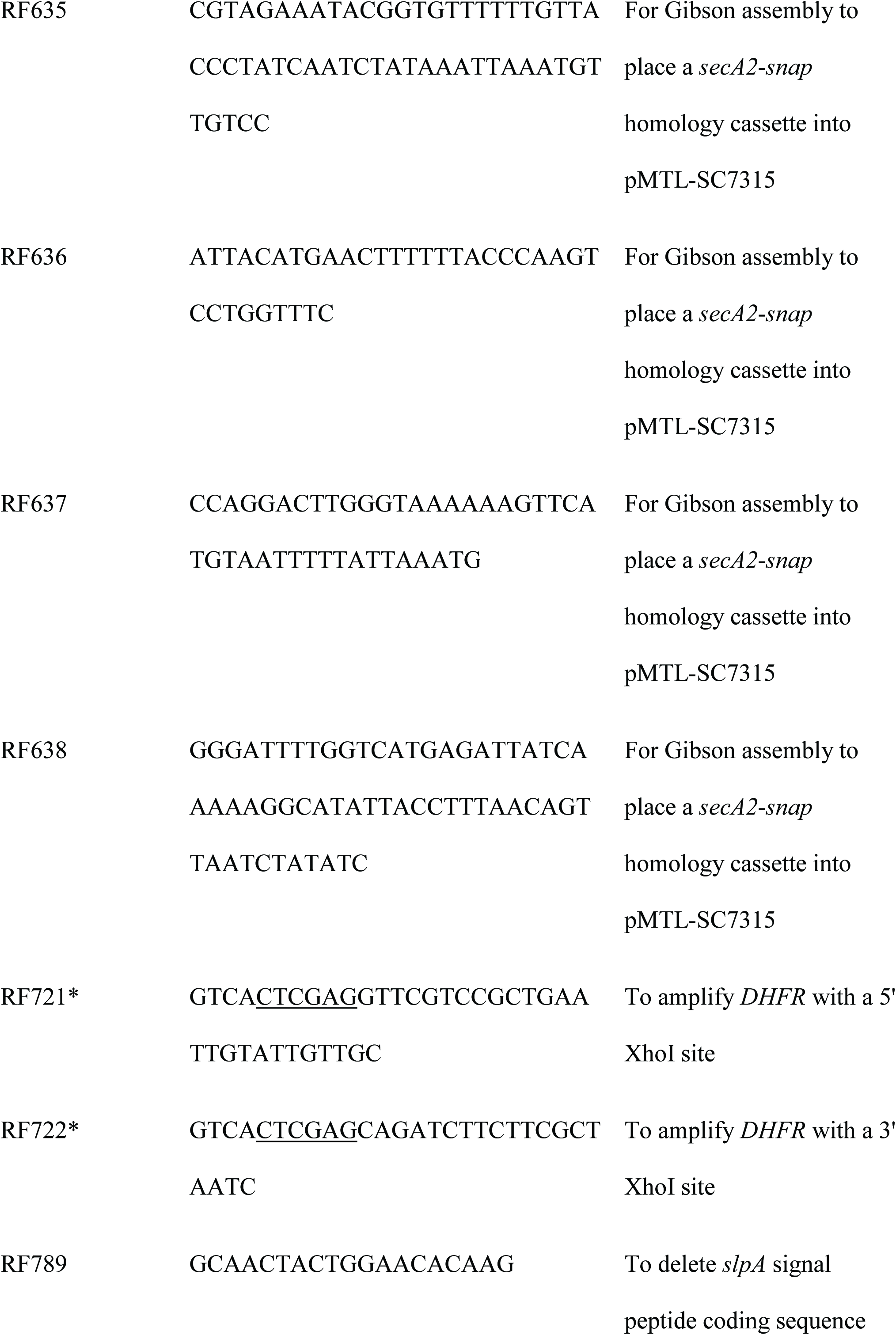
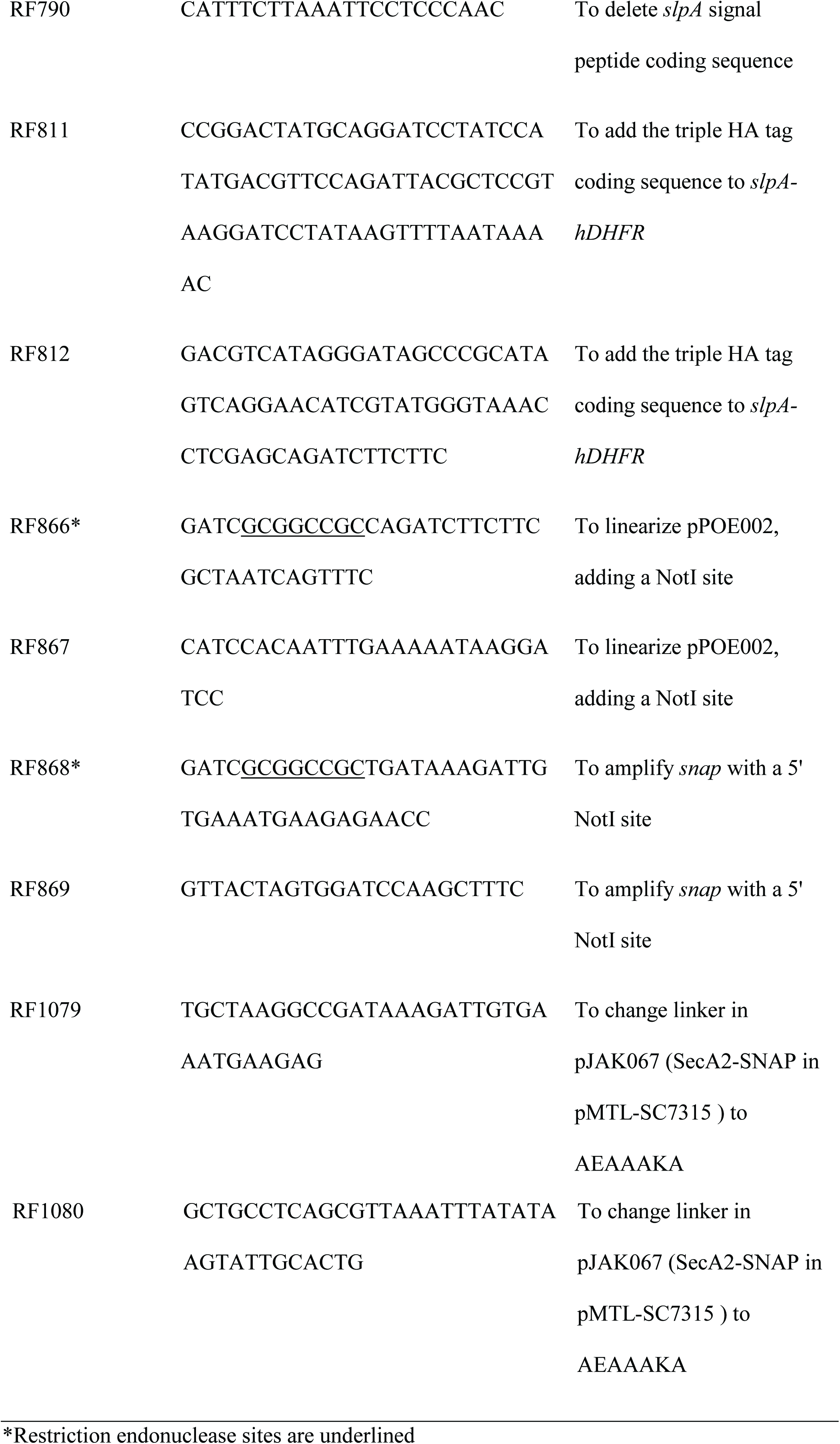
Strains, plasmids and oligonucleotides used in this study

For SlpA-hDHFR-Strep Tag II expression, bacteria were subcultured from overnight cultures to an OD_600nm_ of 0.05, grown for 30 minutes and then supplemented with 200 µM methotrexate. Bacteria were then grown for a further 30 minutes before expression was induced with 20 ng/ml anhydrotetracycline (Atc). Bacteria were grown for a further 3 hours before harvesting at 4,000 xg, for 10 min at 4°C.

### Molecular Biology

Chemically competent *E. coli* were transformed by heat shock using standard methods and plasmids were transferred to *C. difficile* strain 630 by conjugation using the *E. coli* donor strain CA434 (Kirk & Fagan, 2016). Standard techniques were used for PCR, restriction digestion, ligation and Gibson assembly. DNA modifications were performed using Phusion High-Fidelity DNA Polymerase (Thermo Fisher) and Q5 Site-Directed Mutagenesis Kit (NEB) as per manufacturers’ instructions.

### Plasmid Construction

pRFP233 (Kirk et al., 2017) was modified to add the tetracysteine (Tc) tag encoding sequence into *slpA* such that a modified LMW-SLP is produced with FLNCCPGCCMEP added to a surface-exposed loop. The plasmid was linearized by inverse PCR using oligonucleotides RF411 and RF412, deleting 150 bp of the LMW-SLP coding sequence. A synthetic DNA fragment including the deleted 150 bp and 36 bp encoding the Tc tag was inserted by Gibson assembly, yielding plasmid pRPF238.

For the addition of the SNAP tag to SecA2, *secA2* was amplified using RF216 and RF217 from gDNA and digested using SacI/XhoI. *snap* was amplified from pFT46 (Pereira et al., 2013) using RF218 and RF219 and digested with BamHI/XhoI. These fragments were then ligated into SacI/BamHI digested pRPF144 (Fagan & Fairweather, 2011) in a 3-fragment ligation reaction yielding pJAK014. *secA2*-*snap* was excised using SacI/BamHI and ligated into similarly treated pRPF185 yielding pJAK038.

Modification of the *C. difficile* 630 genome was achieved by allele exchange as described previously (Cartman, Kelly, Heeg, Heap, & Minton, 2012). The *snap* coding sequence and the last 1.2 kb of *secA2* was amplified by PCR using RF635 and RF636, using pJAK014 as a template. 1.2 kb downstream of *secA2* was amplified by PCR using RF637 and RF638 using gDNA as a template. pMTL-SC7315 (Cartman et al., 2012) was linearised by PCR using RF311 and RF312. The three fragments were ligated by Gibson assembly yielding pJAK067. To improve the enzymatic activity of the expressed fusion protein, the size of the linker between SecA2 and SNAP was increased. pPOE032 was prepared by inverse PCR of pJAK067 with RF1079 and RF1080 via a site-Directed Mutagenesis Kit (NEB) as per manufacturer’s instructions.

An *slpA*_*630*_*-strep tag II* encoding SacI/BamHI insert was excised from pRPF173 and ligated into a SacI/BamHI digested pRPF185 yielding pPOE005. pPOE005 was modified to add the coding sequence for *hDHFR-myc* within *slpA*_*630*_*-strep tag II*. *hDHFR-myc* was amplified using RF721 and RF722 and ligated into XhoI linearized pPOE005 to give pPOE002. Sequence encoding a 3xHA tag was added to *slpA*_*630*_*-hDHFR-myc* by inverse PCR of pPOE005 with RF811 and RF812 by site-directed mutagenesis, as described earlier, to create pPOE003. The sequence encoding the SlpA_630-_hDHFR-Strep Tag II signal peptide in pPOE002 was removed by inverse PCR site-directed mutagenesis with RF789 and RF790 to create pPOE011. pPOE023 was prepared by ligation of the *slpA*_*630*_ SacI/XhoI insert from pPOE005 into SacI/XhoI digested pJAK038 vector.

To add a SNAP tag to SlpA_630_-hDHFR-Strep Tag II, pPOE002 was linearized by PCR using RF866 and RF867. *snap* was amplified from pJAK038 using RF868 and RF869. These fragments were then NotI/BamHI digested and ligated yielding pJAK085.

### Microscopy

#### SNAP Cell TMR-Star and SNAP-Surface 549 Staining

cells were grown from an OD_600nm_ 0.05 to 0.4 and treated with 250 µM TMR-Star for at least 30 minutes. Transient expression of SlpA-SNAP or SlpA-DHFR-SNAP was induced for 10 minutes with 10 ng/ml Atc before fixation or 20 ng/ml for 1 hour for in-gel fluorescence experiments.

#### HADA Staining

Cells were grown to an OD_600nm_ of approximately 0.1 before the addition of 0.5 mM HADA and continued growth for at least 2 hours to an OD_600nm_ of 0.5-0.6. To chase the HADA staining, cells were harvested at 4,000 x g for 5 minutes, washed once by resuspension in 8 ml reduced TY and finally resuspended in 2x the original volume of reduced TY media before continuing growth for up to 30 minutes. In the case of transient SlpA_R20291_ expression the cells were grown for 25 minutes before inducing the expression with 100 ng/ml Atc for the final 5 minutes before fixation.

Live-cell sample preparation: After the wash with 8 ml TY (as described above), HADA stained cells were resuspended in reduced TY to an OD of approx. 50. 0.5 OD600U of cells were transferred in an anaerobic chamber to a glass bottom petri dish (ibidi) at the interface between the glass coverslip and a 1% agarose pad that covered the entire surface of the dish and had been reduced for at least 3 hours. The petri dish was tightly wrapped in parafilm under anaerobic conditions before immediately been transferred at 37°C to a widefield microscope chamber pre-heated to 37°C for imaging.

#### Fixation

Cells were harvested at 4,000 x g for 5 minutes at 4°C, washed two times in 1 ml ice cold PBS with spins at 8,000 x g for 2 min at 4°C before being fixed with 4% paraformaldehyde in PBS for 30 minutes at room temperature. After fixation, cells were washed three times in PBS. For immunofluorescence the fixed cells were blocked overnight with 3% BSA in PBS at 4°C. Cells were harvested at 8,000 x g for 2 min at 4°C, resuspended in 1:500 Primary antibody (Mouse Anti-027 SlpA_R20291_ LMW-SLP) and incubated at room temperature for 1 hour. Cells were then washed three times in 1 ml 3% BSA in PBS before being resuspended in 1:500 secondary antibody (Goat anti-mouse-Cy5, Thermo Fisher). Cells were incubated for 1 hour at room temperature then washed three times in 3% BSA in PBS before being resuspended in PBS. Washed cells were dried down to glass cover slips and mounted with SlowFade Diamond (Thermo Fisher).

Images were taken on a Nikon Ti eclipse widefield imaging microscope using NIS elements software or a ZEISS LSM 880 with Airyscan using ZEN imaging software. Image J based FiJi was used for image analysis.

### Cell Fractionation

Extracellular protein extraction: Cells were harvested at 4,000 x g for 5 minutes at 4°C. In all the following wash steps bacterial cells we centrifuged at 8,000 x g for 2 minutes. Pellets were washed twice by resuspension in 1 ml ice cold PBS. Cells were treated with 10 µl per OD_600nm_U of extraction buffer (0.2 M Glycine, pH 2.2) and incubated at room temperature for 30 minutes to strip extracellular proteins. Stripped cells were harvested and the supernatant, containing extracellular protein, was taken and neutralized with 0.15 µl 1.5 M Tris pH 8.8 per 1 µl extract. The stripped cells were washed twice in 1 ml ice cold PBS before being frozen at −80C. Cells were thawed and then resuspended in 11.5 µl per OD_600nm_U cell lysis buffer (PBS, 1x protease inhibitor cocktail, 5 mM EDTA, 20 ng/ml DNase, 120 mg/ml purified CD27L endolysin (Mayer, Garefalaki, Spoerl, Narbad, & Meijers, 2011)). Lysis was induced by incubating at 37°C shaking for 30 minutes. Cell membranes were harvested by centrifugation at 20,000 x g for 20 minutes and the soluble intracellular protein fraction retained before the pellet was washed twice with 1 ml PBS. Membranes were solubilized using 11.5 µl per OD_600nm_U solubilization buffer (1x PBS, 1x Protease Arrest, 5 mM EDTA, 20 ng/ml DNase, 1.5% sarkosyl) and agitated by rotating for 1 hour at room temperature. Insoluble material was harvested at 20,000 x g for 5 minutes and the solubilized membrane fraction was taken. Alternatively, for the SNAP tagged SlpA constructs; cells lysates were supplemented with 1.5% sarkosyl, incubated for 1 hour and harvested at 20,000 x g for 5 minutes to create a total cellular extract.

### Protein Gels

Proteins were separated using standard SDS-PAGE techniques on a mini-protein III system (Bio-Rad) before being either; analyzed for in-gel florescence on a ChemiDoc imaging system (Bio-Rad), stained with Coomassie or transferred to nitrocellulose membranes using a semi-dry blotter (Bio-Rad) for western blot analysis. Band intensities were measured using Image Lab Software (Bio-Rad).

### Statistics

Statistics were performed in Origin using one-way analysis of variance (ANOVA), a difference with p ≤ 0.05 was considered significant.

## Supporting information

## Acknowledgements

We would like to thank Darren Robinson and Christa Walther at The Wolfson Light Microscopy Facility at the University of Sheffield for their help with microscopy. We would also like to thank Aimee Shen for helpful discussion and Neil Fairweather for feedback on a previous version of this manuscript.

This work was supported by the Medical Research Council (grant number MR/N000900/1) and the Wellcome Trust (grant number 204877/Z/16/Z).

## Supplemental Figure Legends

**Figure 2-figure supplement 1:**
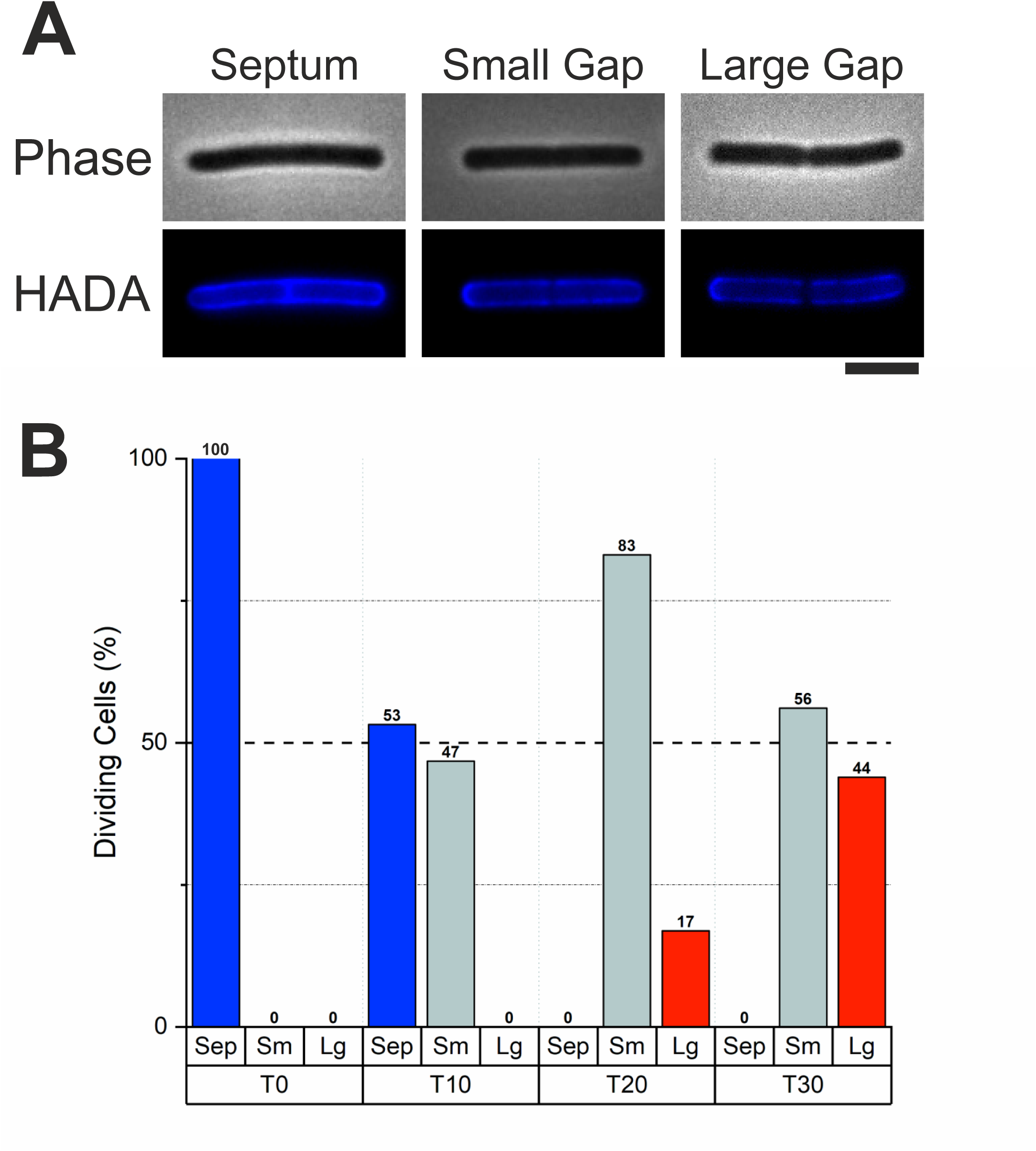
*C. difficile* 630 HADA staining chase time course. **A:** Widefield microscopy displaying phase contrast (upper panels) and fluorescence (lower panels) of HADA stained *C. difficile* 630 cells chased for 0, 10, 20 or 30 min without HADA (as described in Methods). HADA staining at the center of a dividing cell can be characterized as: septum stained (left) or patches of reduced HADA staining being smaller than 360 nm in length (middle) or larger (right). Scale bar indicates 3 µm. **B:** Graph displaying the population distribution of dividing *C. difficile* 630 cells characterized for HADA staining in widefield microscopy (as in Figure 2S1A) when chased for HADA for 0, 10, 20 or 30 minutes (T0, n=56; T10, n=62; T20, n=71; T30, n=107). The percentage of the total counted population are displayed above each bar.

**Figure 2-figure supplement 2:**
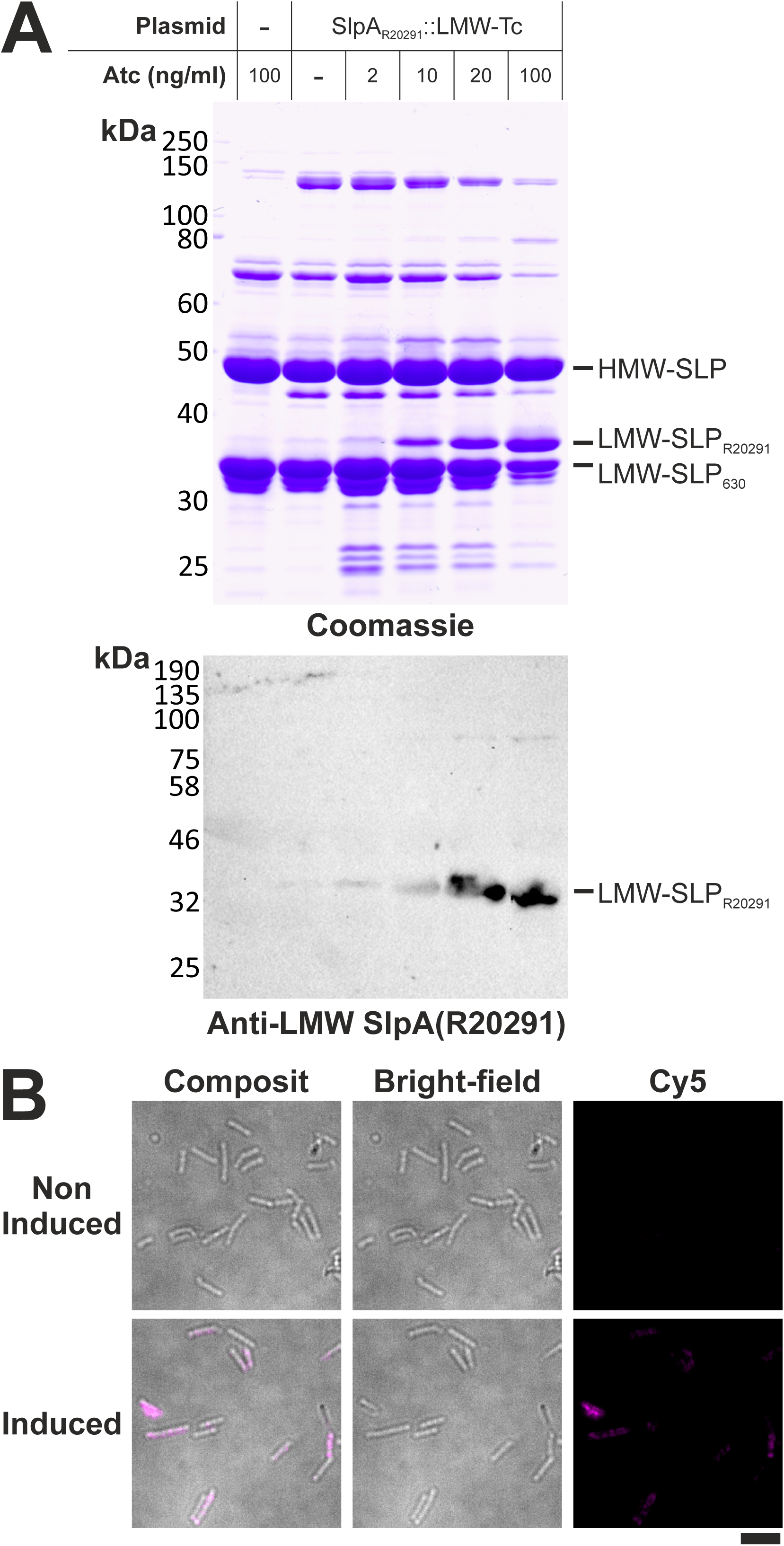
Antibody specificity for SlpA_R20291_ LMW-SLP. **A:** Top panel: Coomassie stain of SDS-PAGE separated extracellular extracts from *C. difficile* 630 cells grown for three hours with the indicated amount of anhydrotetracycline (Atc) to induce protein expression. Lower panel: Western immunoblot to detect SlpA_R20291_ LMW-SLP in the same extracellular extracts. **B:** Widefield microscopy of *C. difficile* 630 cells with pRPF238 (encoding SlpA_R20291_::LMW-Tetracysteine. The tetracysteine tag was used during other labelling experiments that were unsuccessful (data not shown)) induced (bottom panels) or not induced (top panels) with 100 ng/ml Atc for 5 minutes. Surface SlpA_R20291_ was immunolabeled with Cy5 (magenta). Scale bar indicates 6 µm.

**Figure 2-figure supplement 3:**
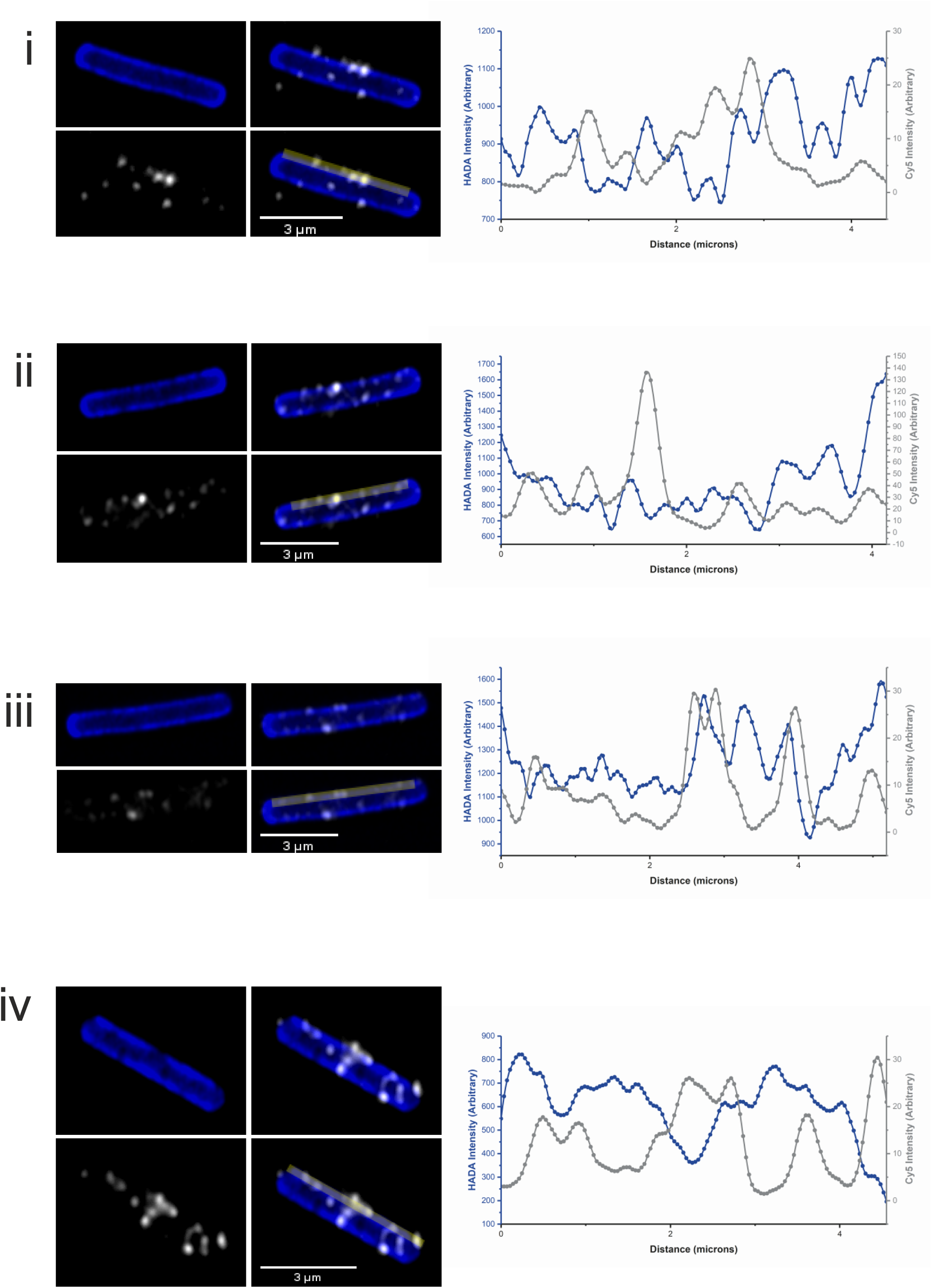
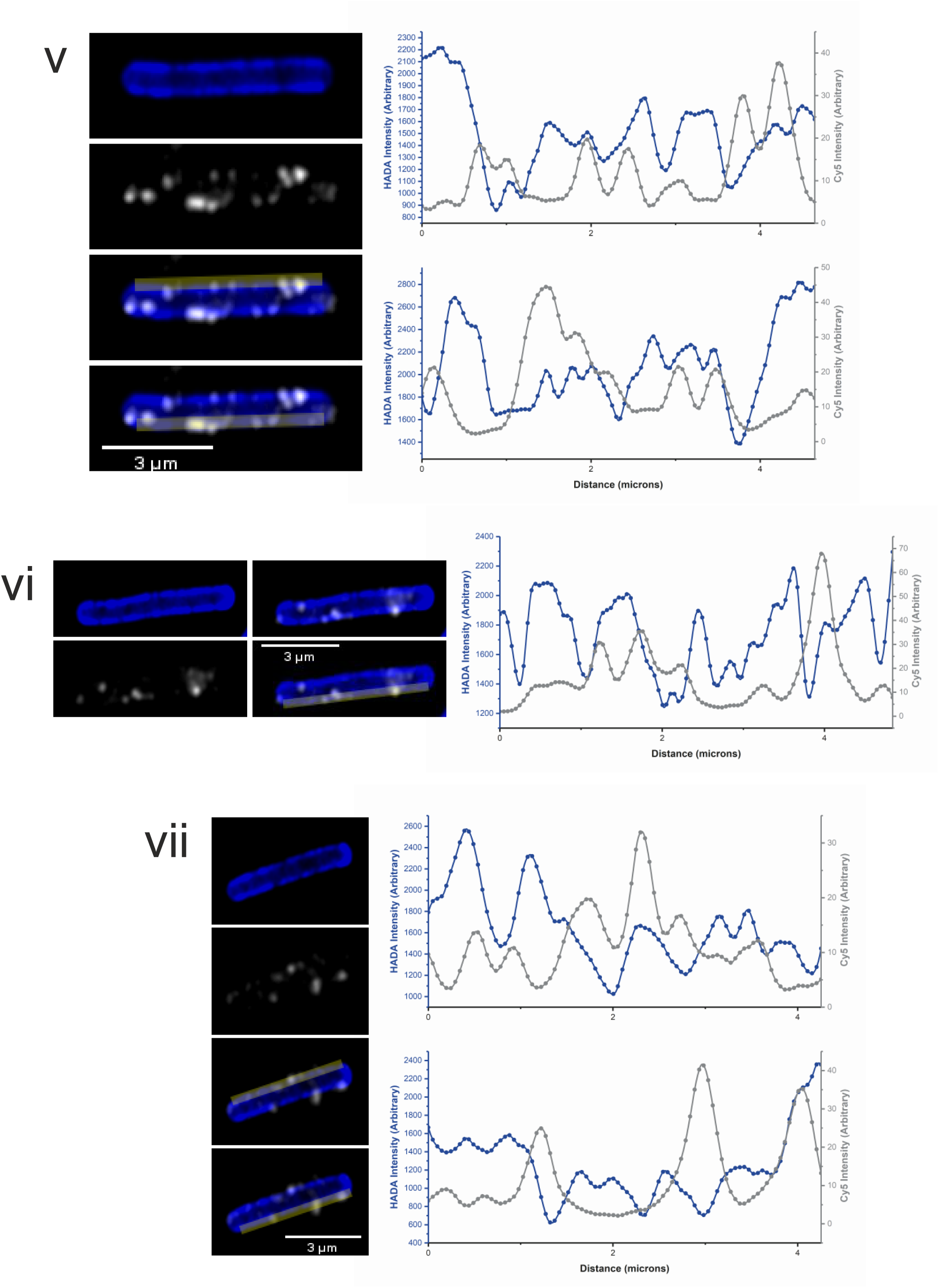
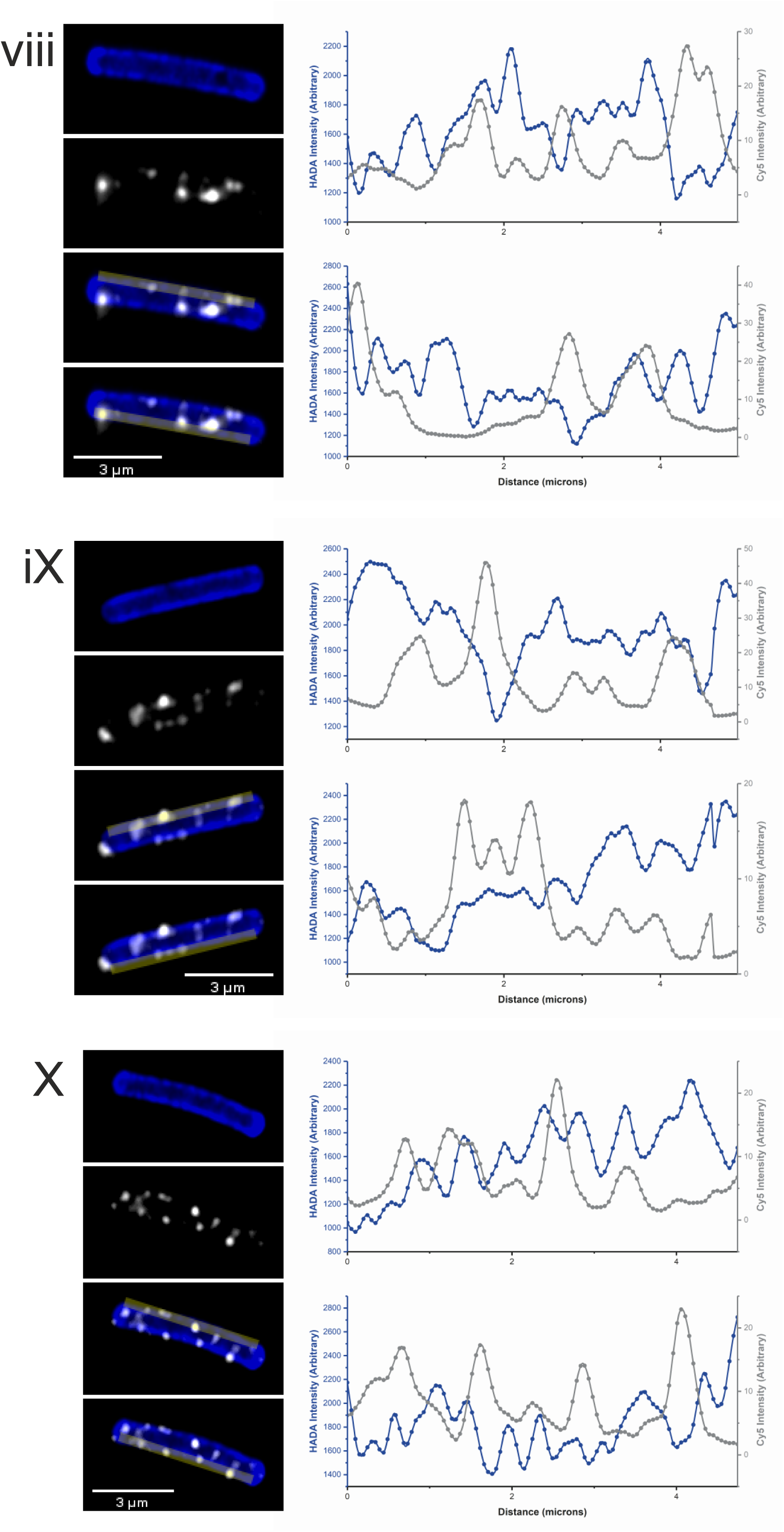
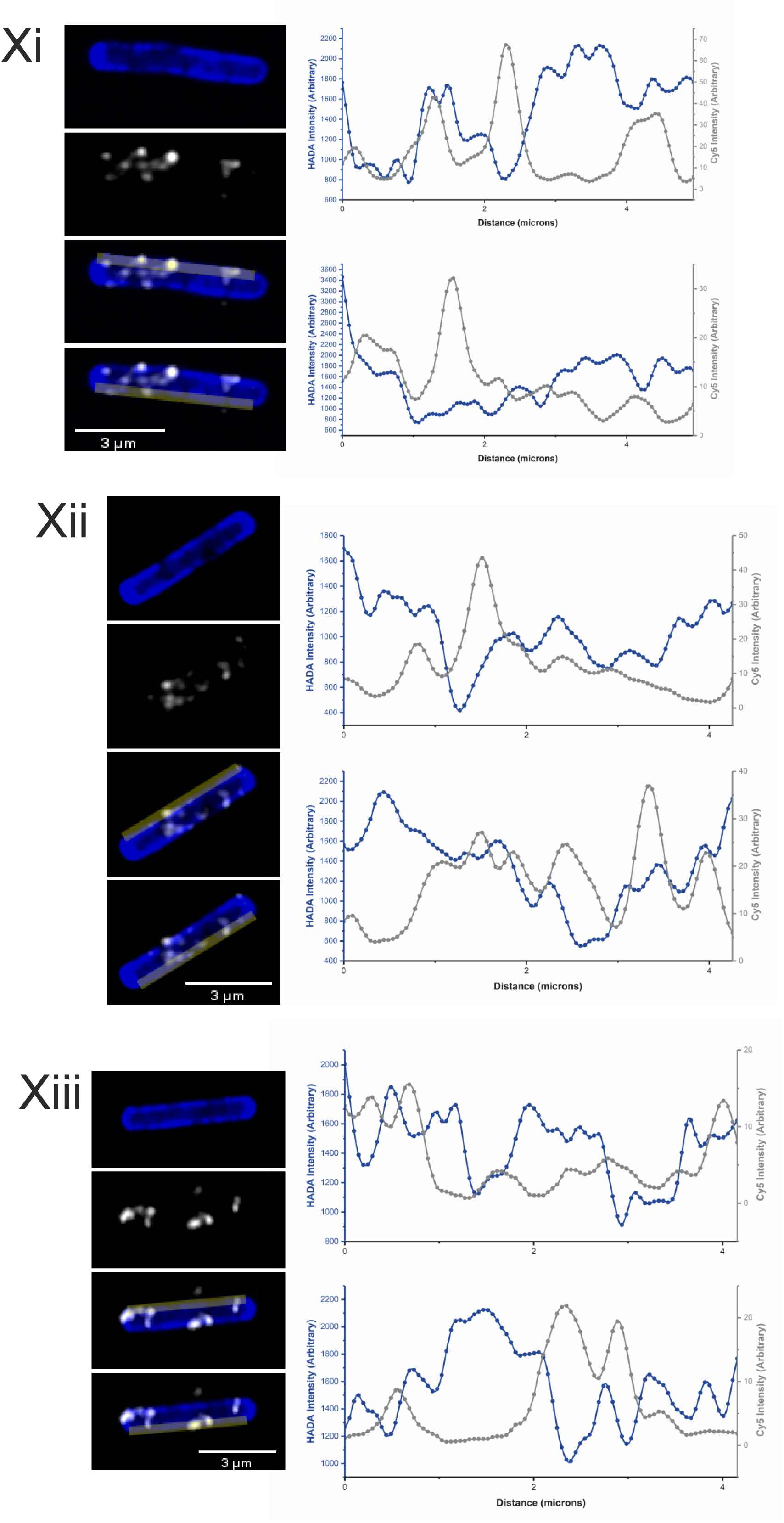
Further images of new surface S-layer. Representative examples of airyscan confocal images displaying *C. difficile* 630 cells prepared as in Figure 2 with HADA label peptidoglycan cell wall (Blue), new SlpA_R20291_ immunolabeled with Cy5 (White) and yellow bar regions used for intensity plot graphs. Intensity plot graphs display HADA (Blue) and Cy5 (Grey) signal with upper and lower graphs corresponding to the higher and lower cell region marked with yellow bars in the left panels, respectively.

**Figure 4-figure supplement 1:**
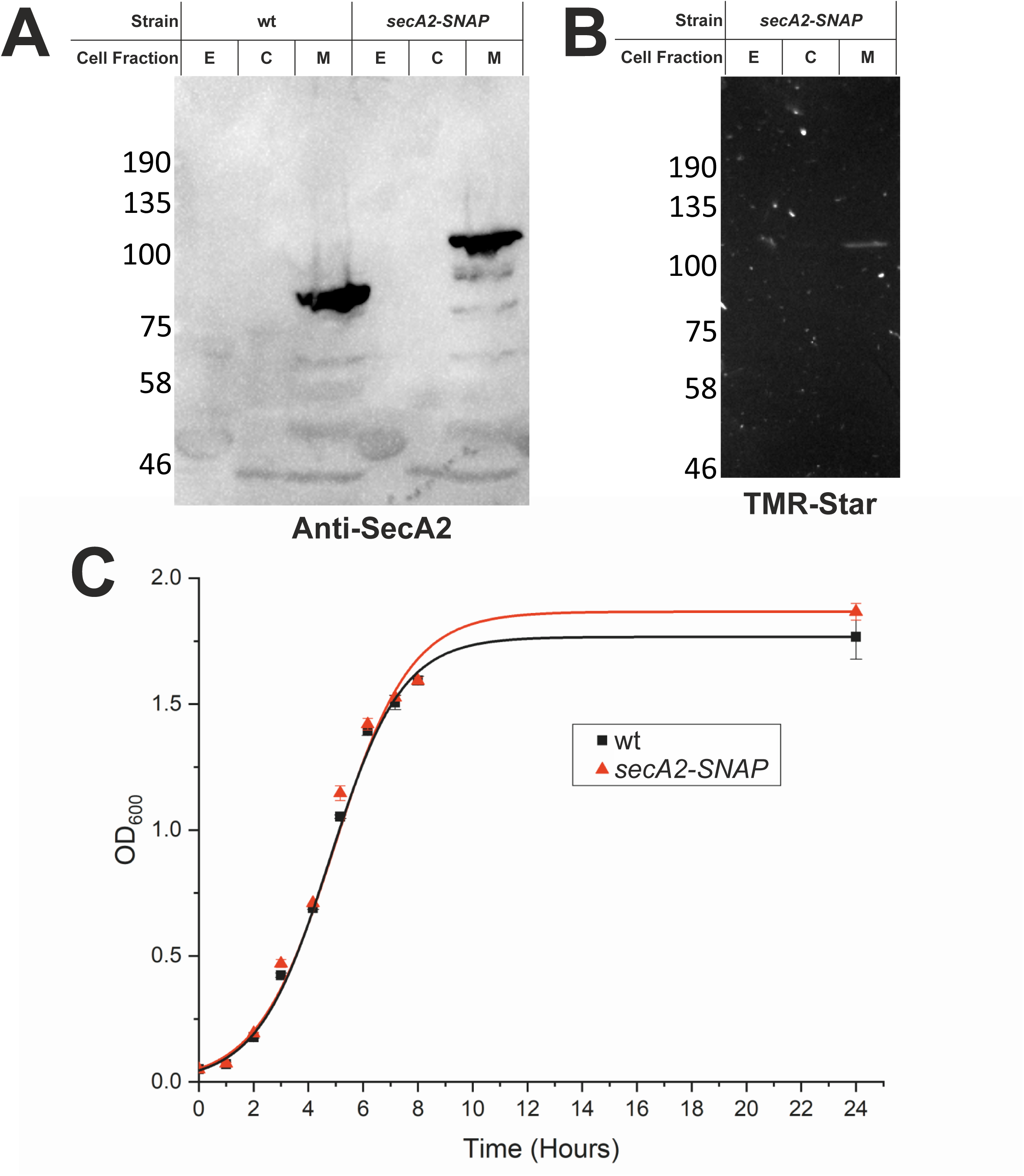
SecA2-SNAP is functional in *C. difficile.* **A:** Western immunoblot showing the distribution of SecA2 in extracellular (E), cytosolic (C) and membrane (M) fractions from wild-type *C. difficile* 630 or cells expressing a genomic copy of a *secA2-SNAP* fusion, loaded at the same OD_600_U. **B:** In-gel fluorescence of SecA2-SNAP-TMR-Star from cell extracts expressing SecA2-SNAP (labelled as in A). **C:** Growth curves of wild-type *C. difficile* 630 (wt) or 630 *secA2-SNAP.* Following inoculation at an OD_600_ of 0.05, growth was followed by measuring OD_600_ hourly. Shown are the mean and standard error of duplicate cultures.

**Figure 5-figure supplement 1:**
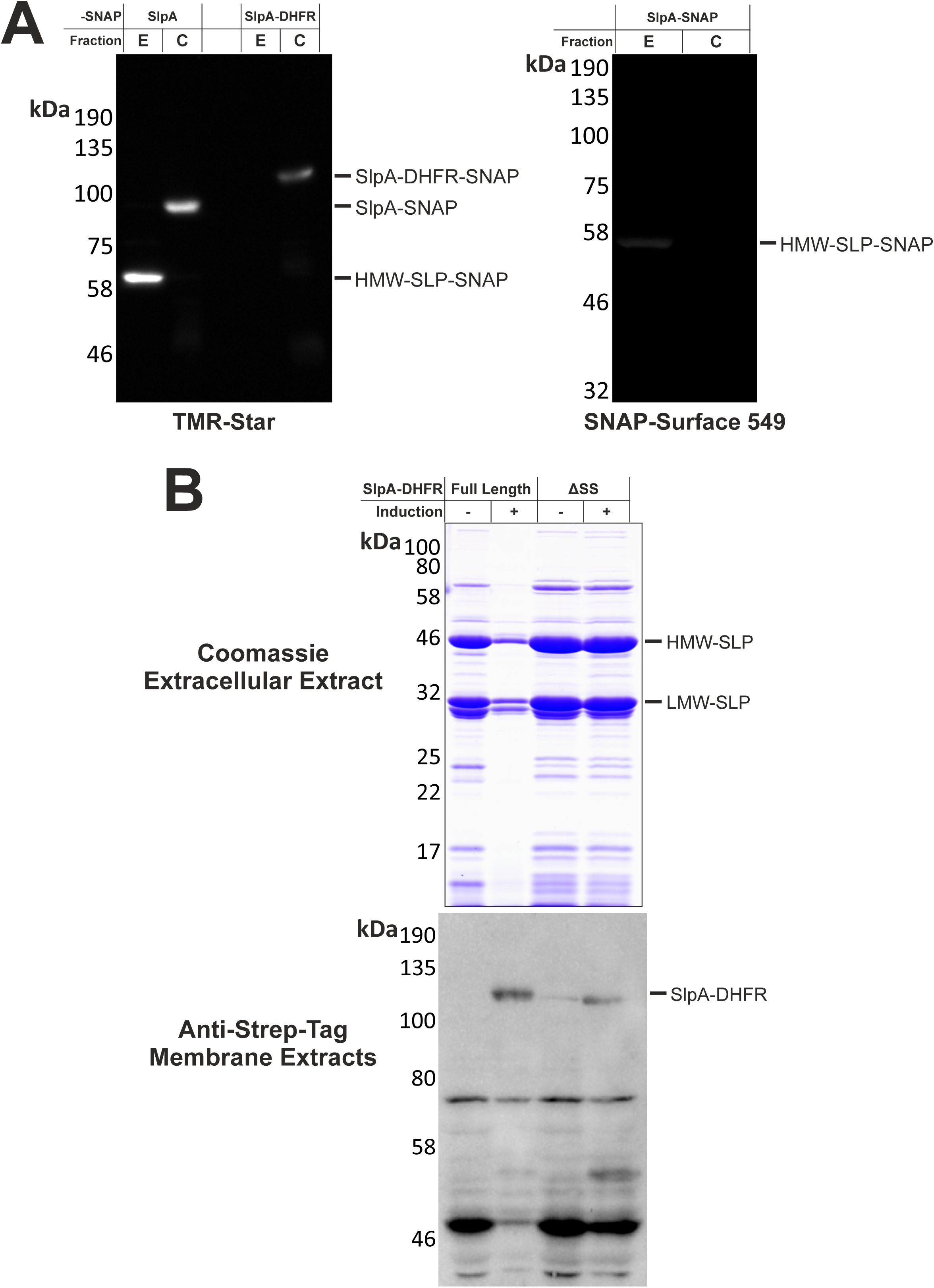
SlpA-SNAP, SlpA-DHFR-SNAP, SlpA-DHFR expression and localization. **A:** SDS PAGE in-gel fluorescence displaying SNAP-TMR-Star signal (left) or SNAP-Surface 549 (right) from extracellular (E) or cellular (C) *C. difficile* 630 extracts expressing SlpA-SNAP or SlpA-DHFR-SNAP. **B:** SDS PAGE analysis of extracellular extracts stained with coomassie (upper panel) or membrane fractions analyzed by Western immunoblot with an anti-strep-tag antibody (lower panel) from *C. difficile* 630 cells expressing strep tagged full length SlpA-DHFR or SlpA-DHFR lacking a signal sequence (ΔSS). Protein expression was induced with 20 ng/ml Atc for 180 min as indicated.

**Figure 5-figure supplement 2:**
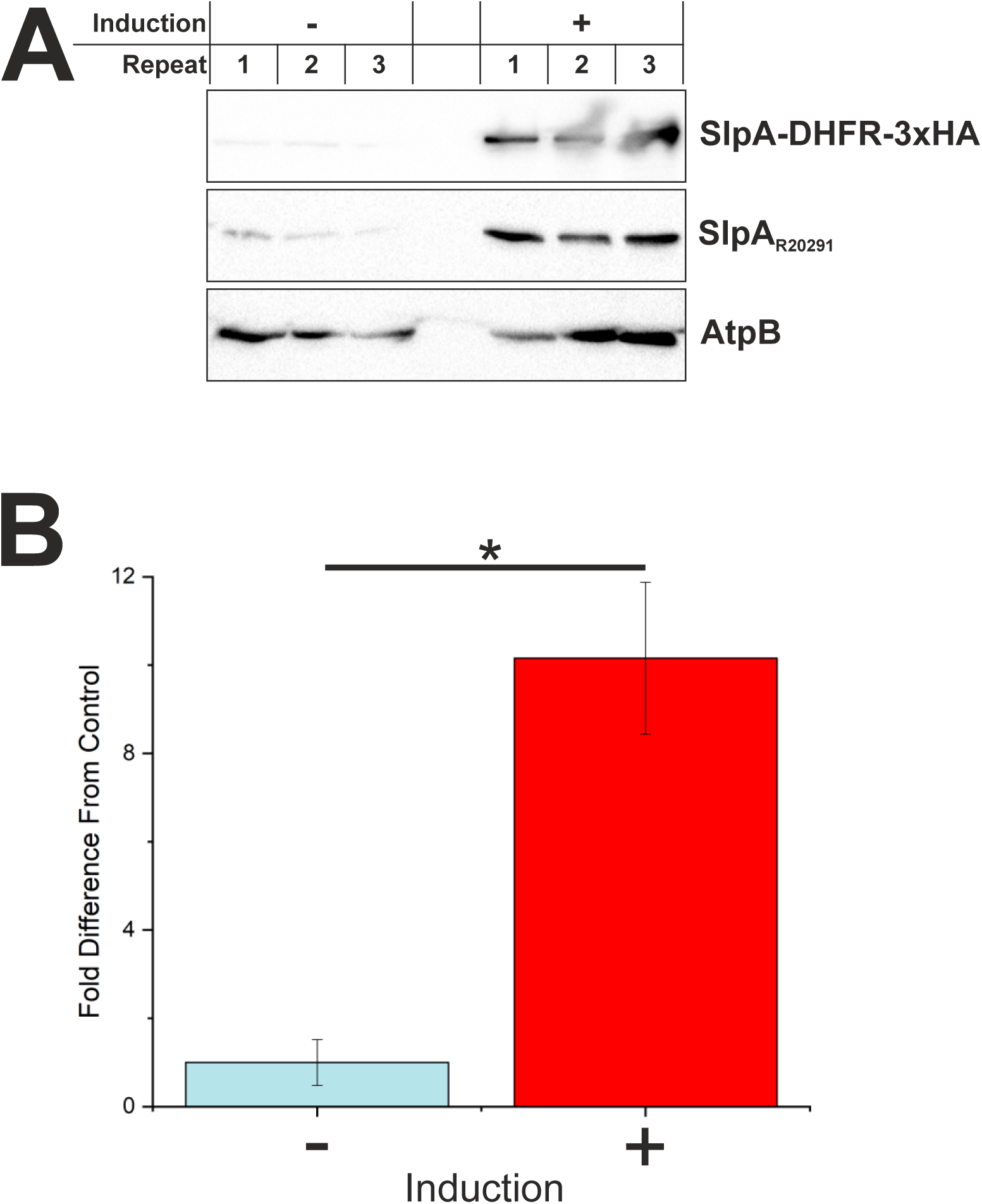
Characterization of a SlpA-DHFR fusion protein. **A:** Western immunoblot analysis of *C. difficile* R20291 expressing an SlpA-DHFR-3xHA fusion. Protein expression was induced (+) with 20 ng/ml Atc for 1 hour and intracellular cell extracts (membrane and cytosol) were analyzed by SDS PAGE followed by Western immunoblot using an anti-HA antibody to show expression of SlpA-DHFR-3xHA (top panel), anti-SlpA_R20291_ to visualize accumulation of native SlpA precursor in the cytosol (middle panel) and anti-AtpB as a membrane protein and loading control (bottom panel). Samples from triplicate cultures are shown. **B:** Quantification of average fold change of intracellular SlpA_R20291_ precursor band intensity from A. Native SlpA_R20291_ secretion is blocked by expression of SlpA-DHFR-3xHA. Asterisk indicates a significant difference (p ≤ 0.05).

**Figure 5-figure supplement 3:**
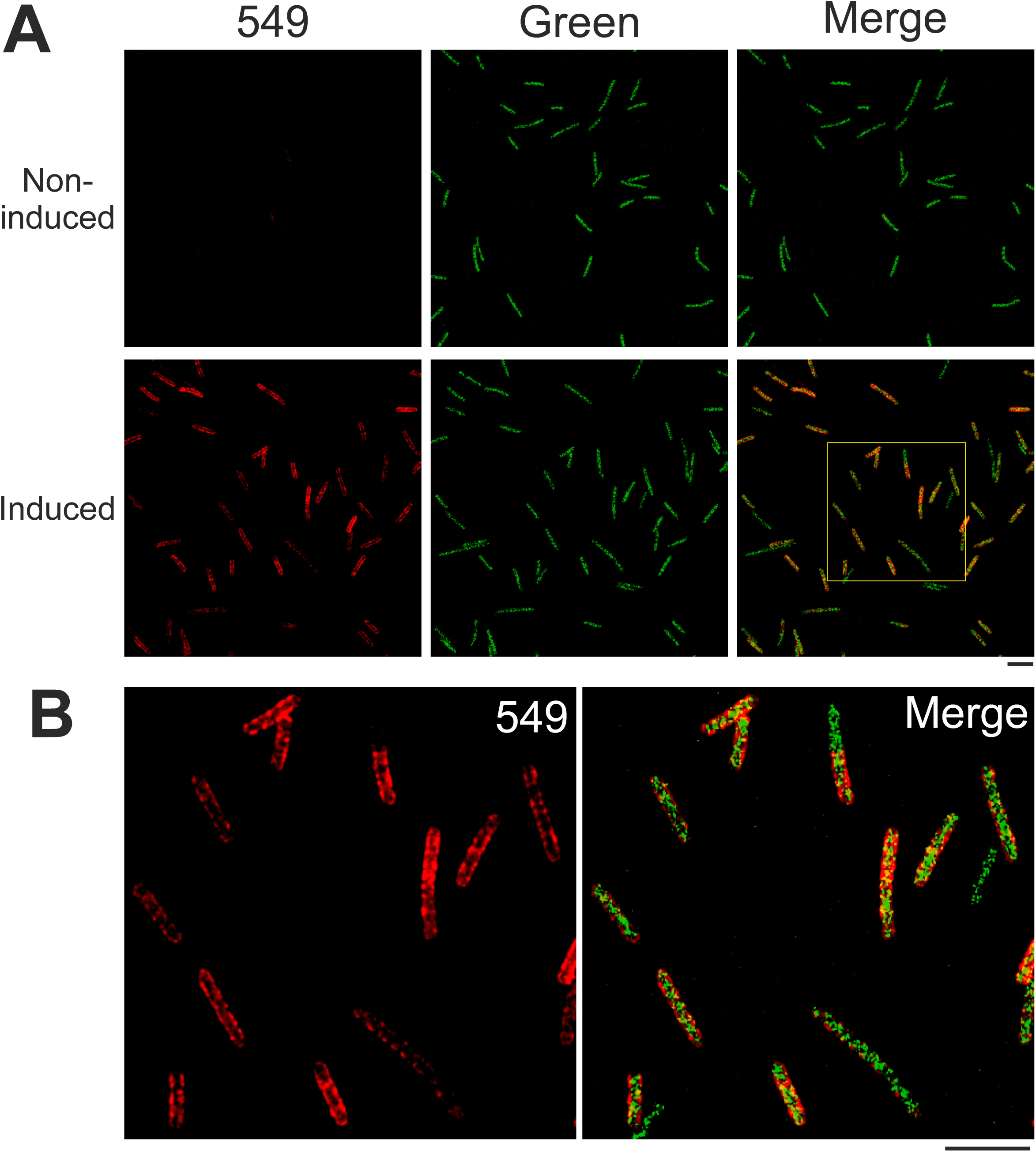
SNAP-Surface 549 Stained HMW-SlpA-SNAP. **A:** Airyscan confocal images of *C. difficile* 630 cells stained with SNAP-Surface 549 and induced or not induced for SlpA-SNAP expression. Surface 549 signal (left panels), green autofluorescence from *C. difficile* 630 cells (middle panels) and merged (right panels). Area taken for zoomed image depicted by a yellow square. Scale bar indicates 6 µm. **B:** Zoomed area (from **A**) of HMW-SlpA-SNAP-Surface 549 signal (left panel) and autofluorescence merged image (right panel). Scale bar indicates 6 µm.

### Video 1: Live-cell imaging of *C. difficile* HADA Chase

Live-cell widefield microscopy of HADA fluorescent signal (top panel) and phase contrast (bottom panel) from *C. difficile* 630 cells chased for HADA staining, with a cell undergoing the final stages of cell division. 22 frames at ∼3 minutes per frame, scale bar indicates 3 µm.

