## Supplementary figures and images for "Spatial organization of *Clostridium difficile* S-layer biogenesis"

### Supplementary file 1

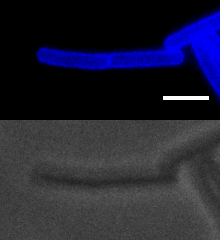
